# In *Hyphomicrobium denitrificans* two related sulfane-sulfur responsive transcriptional repressors regulate thiosulfate oxidation and have a deep impact on nitrate respiration and anaerobic biosyntheses

**DOI:** 10.1101/2025.02.17.638619

**Authors:** Jingjing Li, Nora E. Schmitte, Kaya Törkel, Christiane Dahl

**Affiliations:** Institut für Mikrobiologie & Biotechnologie, Rheinische Friedrich-Wilhelms-Universität Bonn, Meckenheimer Allee 168, 53115 Bonn, Germany

**Keywords:** *Hyphomicrobium denitrificans*, thiosulfate oxidation, transcriptional regulation, denitrification, PQQ, iron acquisition, ubiquinone biosynthesis

## Abstract

Bacteria have evolved multiple strategies to sense and respond to the availability of inorganic reduced sulfur compounds such as thiosulfate. In *Hyphomicrobium denitrificans*, an obligately chemoorganoheterotrophic Alphaproteobacterium, the use of thiosulfate as a supplemental electron donor is regulated by two homologous sulfane-sulfur-responsive ArsR-type transcriptional repressors, sHdrR and SoxR. Here, we provide information on the distribution and phylogeny of sHdrR, the relevance of its two conserved cysteines i*n vivo*, and identify the genes controlled by SoxR and sHdrR not only by targeted qRT-PCR but also by global RNA-Seq-based analyses of regulator-deficient mutant strains.The absence of sHdrR and SoxR affected 165 and 170 genes, respectively, with 138 genes overlapping. SoxR affects the *sox* genes for periplasmic thiosulfate oxidation and sulfane sulfur import into the cytoplasm, as well as the *lip-shdr-lbpA* genes encoding the cytoplasmic enzymes essential for sulfite formation. sHdrR affects only a subset of these genes. The transcription of *sox* genes remains unaltered in its absence. sHdrR and SoxR act cooperatively, possibly involving heterodimer formation, and their activity also involves interaction with other transcriptional regulators. Most importantly, sHdrR/SoxR regulation extends far beyond sulfur oxidation and deeply affects anaerobic metabolism, particularly denitrification in *H. denitrificans*.

## INTRODUCTION

Most dissimilatory sulfur oxidizing prokaryotes oxidize thiosulfate (S_2_O_3_^2-^), an inorganic sulfur compound of intermediary redox state. Accordingly, microbial thiosulfate oxidation has a major impact on global sulfur cycling. Thiosulfate oxidation under aerobic conditions is very wide-spread, particularly in marine environments (Podgorsek and Imhoff, 1999; Marshall and Morris, 2013; Watsuji et al., 2016). Anaerobic thiosulfate oxidizers have also been reported from a large range of environments, including anoxic marine basins (Menezes et al., 2020), oceanic oxygen minimum zones (Callbeck et al., 2021), and hydrothermal vents (Teske et al., 2000). In fact, it is likely that anaerobic oxidation, primarily through nitrate and nitrite reduction, is responsible for much of the thiosulfate removal in the marine environment (Ding et al., 2023). Thiosulfate-dependent denitrification to N_2_ is best known for obligately autotrophic species such as *Thiobacillus denitrificans*, *Thiomicrospira denitrificans* or *Sulfurovum lithotrophicum* (Inagaki et al., 2004), has been found to be important in marine chemosynthetic symbioses (Paredes et al., 2021), and has also been reported for the facultatively autotrophic *Paracoccus pantotrophus* (Robertson and Kuenen, 1983).

Sulfur oxidizers include not only autotrophic prokaryotes from various groups, but also a large number of obligately organoheterotrophic bacteria that oxidize thiosulfate as an additional electron donor and are widely distributed in soil and natural waters (Trudinger, 1967; Tuttle and Jannasch, 1972; Sorokin et al., 1999; Ding et al., 2023). These include species of the genus *Hyphomicrobium*, Alphaproteobacteria that can be isolated from virtually any freshwater or soil sample, where they can constitute up to 0.2% of the total bacteria (Hirsch and Conti, 1964; Gliesche et al., 2015; Li et al., 2023b). They are also prevalent in temporary puddles and in activated sludge, even under anaerobic conditions. Denitrifying hyphomicrobia such as *H. denitrificans*, are of particular interest because of the need to remove nitrate in drinking water and sewage treatment plants, where these organisms are indeed highly abundant (Holm 1996). *Hyphomicrobium* spp. are typically restricted to C_1_ and C_2_ compounds as carbon sources and are commonly identified as major players in denitrification systems supplied with methanol (Martineau et al., 2015). While enzymes required for denitrification have been purified and characterized from *H. denitrificans* (Deligeer et al., 2002; Yamaguchi et al., 2003; Yamaguchi et al., 2004), and genetic and physiological aspects of its denitrification are beginning to emerge (Martineau et al., 2015), these studies have not yet included any aspects of oxidative sulfur metabolism.

In *H. denitrificans* thiosulfate oxidation commences in the periplasm. Here, two thiosulfate molecules can be oxidatively linked to form the dead-end product tetrathionate, a reaction catalyzed by thiosulfate dehydrogenase (TsdA) (Koch and Dahl, 2018; Li et al., 2023b). Alternatively, thiosulfate can be completely oxidized to sulfate. This pathway is preferred at lower substrate concentrations (<2.5 mM) and involves the periplasmic SoxYZ carrier protein to which thiosulfate is oxidatively bound by the action of the *c*-type cytochrome SoxXA (Li et al., 2023b). Sulfate is then hydrolyzed off by SoxB and the sulfane sulfur remaining on SoxYZ is transferred to the cytoplasm via the membrane transporter SoxT1A (Li et al., 2024). Once inside, the sulfur is delivered through a cascade of sulfur transfer reactions to the sulfur-oxidizing heterodisulfide-reductase-like enzyme complex, sHdr (Tanabe et al., 2024), which releases sulfite as a product. Sulfite is excreted and, in the absence of efficient sulfite-oxidizing enyzmes in *H. denitrificans*, chemically oxidized to sulfate in the presence of oxygen or may be transformed by other community members under environmental conditions (Li et al., 2023b; Li et al., 2024).

In the environment, *H. denitrificans* must not only cope with constantly changing concentrations of respiratory electron acceptors (oxygen, nitrate), but may also encounter varying concentrations of reduced sulfur compounds such as thiosulfate. To make the most of these additional electron sources, the organism must have strategies for sensing their presence. Indeed, we have recently identified two distinct but closely related ArsR-SmtB family transcriptional repressors, SoxR and sHdrR, that are responsible for the transcriptional regulation of genes encoding Sox, sHdr and associated proteins in *H. denitrificans* (Li et al., 2023b; Li et al., 2023a; Li et al., 2024). SoxR has been extensively characterized at the molecular level. Its sensing mechanism involves the formation of a sulfane sulfur bridge between two conserved cysteine residues in the presence of thiosulfate. In the sulfur-bridged form, the repressor can no longer bind to its operator region(s) on the DNA and transcription is released (Li et al., 2023a). Much less is known about sHdrR. It appears to regulate the sHdr system by modulating sHdrA levels, as seen in Western blot experiments, and to bind to the DNA region upstream of its own gene (Li et al., 2023b). Whether and how the two repressors interact in regulating the overall sulfur oxidation process and possibly other parts of hyphomicrobial energy conservation has not been studied in detail.

Here, we address these knowledge gaps by first providing insights into the relationship between SoxR and sHdrR, the distribution and phylogeny of sHdrR, and the relevance of the conserved sHdrR cysteines i*n vivo*, and then collecting information on the SoxR and sHdrR regulons not only by targeted qRT-PCR but also by a global RNA-Seq-based analysis of regulator-deficient mutant strains, the latter revealing a profound effect on anaerobic metabolism, in particular denitrification.

## EXPERIMENTAL PROCEDURES

### Bacterial strains, plasmids, primers, and growth conditions

*H. denitrificans* strains were cultured in minimal medium kept at pH 7.2 with 100 mM 3-(*N*-Morpholino)propanesulfonic acid (MOPS) buffer as previously described (Koch and Dahl, 2018). Media contained 24.4 mM methanol. Thiosulfate was added as needed. *Escherichia coli* strains were grown on complex lysogeny broth (LB) medium (Bertani, 2004) under aerobic conditions at 37°C unless otherwise indicated. *Escherichia coli*. BL21 (DE3) was used for recombinant protein production. *E. coli* strains 10-beta and DH5α were used for molecular cloning. Antibiotics for *E. coli* and *H. denitrificans* were used at the following concentrations (in μg ml^−1^): ampicillin, 100; kanamycin, 50; streptomycin, 200; chloramphenicol, 25. Supplementary Table 1 lists the bacterial strains, and plasmids that were used for this study.

### Recombinant DNA techniques

Restriction enzymes, T4 ligase and Q5 polymerase were obtained from New England Biolabs (Ipswich, UK) and used according to the manusfacturer’s instructions. Oligonucleotides were obtained from Eurofins Genomics Germany GmbH (Ebersberg, Germany). Standard techniques for DNA manipulation and cloning were used unless otherwise indicated (Ausubel et al., 1997). Plasmid DNA from *E. coli* was purified using the GenJET Plasmid Miniprep kit (Thermo Scientific, Waltham, USA). Chromosomal DNA from *H. denitrificans* strains was prepared using the Simplex Easy DNA Extract Kit (GEN-IAL GmbH, Troisdorf, Germany). DNA fragments were extracted from agarose gels using the GeneJET Gel Extraction Kit (Thermo Scientific, Waltham, USA).

### Overproduction and purification of recombinant truncated sHdrR

A truncated version of the *H. denitrificans shdrR* gene not encoding the first 25 amino acids of the wildtype protein was amplified from *H. denitrificans* genomic DNA using primers FrpET22b-sHdrR-trun-NdeI and Rev-pET22b-0682-NotI and (Supplementary Table 1) and cloned between the NdeI and NotI sites of pET22b (+), resulting in pET22bHdsHdrR-trunc. Recombinant sHdrR was overproduced in *E. coli* BL21(DE3). The cells were grown at 37°C in 200 ml LB medium containing ampicillin up to an OD600 of 0.6. Expression of *shdrR-trunc* was induced by adding 0.5 mM IPTG. IPTG-induced *E. coli* cells were grown over night at 20°C. The carboxy-terminally His-tagged protein was purified by affinity chromatography on Ni^2+^-NTA using the same conditions as described for the full length protein (Li et al., 2023b)

### Construction of *H. denitrificans* mutant strains

The suicide plasmid pk18*mobsacB* (Schäfer et al., 1994) and the tetracycline cassette from pHP45Ω-Tc (Fellay et al., 1987) were used for reverse genetics in *H. denitrificans*. Derivatives were constructed using on the basis of previously published procedures (Cao et al., 2018; Koch and Dahl, 2018). For chromosomal complementation of the *H. denitrificans* Δ*tsdA* Δ*shdrR* strain, the *shdrR* gene was amplified together with upstream and downstream regions using primers Fwd_deltaHden0682_BamHI and Rev_deltaHden0682_XbaI and cloned into the XbaI/BamHI sites of pk18*mobsacB*. For chromosomal integration of the genes encoding sHdrR Cys^50^Ser, sHdrR Cys^116^Ser and sHdrR Cys^50^Ser Cys^116^Ser, the modified genes and upstream and downstream sequences were amplified by SOE PCR using the appropriate primers listed in Supplementary Table 1. Finally, the tetracycline resistance cassette from pHP45ΩTc was inserted into each of the plasmids using SmaI. The final constructs were electroporated into *H. denitrificans* Δ*tsdA* Δ*shdrR* and transformants were selected using previously published procedures(Cao et al., 2018; Koch and Dahl, 2018). Single crossover recombinants were Cm^r^ and Tc^r^. Double crossover recombinants were Tc^s^ and survived in the presence of sucrose due to loss of both, the vector-encoded levansucrase (SacB) and the tetracyclin resistance gene. The genotype of the *H. denitrificans* strains generated in this study were confirmed by PCR and sequencing.

### Characterization of phenotypes, quantification of sulfur compounds and biomass content

Growth experiments with *H. denitrificans* were run in medium with 24.4 mM methanol in Erlenmeyer as described earlier(Li et al., 2023b). 2 mM thiosulfate were added when needed. Biomass content, thiosulfate and sulfite concentrations were determined by previously described methods(Dahl, 1996; Li et al., 2023b). All growth experiments were repeated three to five times. Representative experiments with two biological replicates for each strain are shown. All quantifications are based on at least three technical replicates.

### Electrophoretic mobility shift assays (EMSA)

The binding reaction mixtures for EMSA assays (15 μl final volume), contained purified truncated sHdrR protein in various concentrations (up to 400 nM), 2 μl 50 % glycerol and 1.5 μl 10 × binding buffer (100 mM Tris-HCl, 500 mM KCl, 10 mM DTT, 5 % glycerol, pH 8.0). Reaction mixtures were pre-incubated for 20 min at room temperature followed by a further 30 min incubation at 30°C after adding the DNA probe to a final concentration of 17 nM. DNA probes were prepared using the primers listed in Supplementary Table S1. The DNA probes were the same as described in (Li et al., 2023a). After pre-running 6% native polyacrylamide gels at 100 V for 1 h at 4 °C with 0.25 × TBE buffer (25 mM Tris-borate, 0.5 mM EDTA), they were loaded with the reaction mixtures. 0.25 × TBE with 0.5 % glycerol was used as running buffer. The gels were run at 180 V for 1h at 4 °C. Gels were subsequently stained for 20 min with SYBR green I. The bands corresponding to sHdrR-bound and free DNAs were visualized with a ChemiDoc Imaging System (BioRad).

### Expression studies based on RT-qPCR

Total RNA of the relevant *H. denitrificans* strains was isolated from cells harvested in mid-log phase according to an established procedure (Li et al., 2023a). RNA samples of 100 ng were used for RT-qPCR analysis which was performed with the primers listed in Supplementary Table 1 following the method described in (Li et al., 2023a).

### Genome-wide transcriptomic analysis of *H. denitrificans* strains Δ*tsdA* Δ*shdrR* and Δ*tsdA* Δ*soxR* in the absence of thiosulfate

For transcriptome sequencing (RNA-Seq), *H. denitrificans* strains Δ*tsdA* Δ*shdrR* and Δ*tsdA* Δ*soxR* were cultured in 50 ml minimal medium containing 24.4 mM methanol in 200 ml Erlenmeyer flasks at 30°C with shaking at 200 rpm to early log phase. Cells from 20 ml culture were harvested and flash frozen in liquid N_2_ and stored at −70°C. As described in (Li et al., 2024), the RNA was purified with the FastGene RNA Premium Kit (NIPPON Genetics Europe, Düren, Germany) according to the manufacturer’s instructions and with introduction of a modified cell lysis step by bead beating. RNA quality was checked on 1% agarose gels and its concentration was measured using NanoPhotometer NP80 (IMPLEN, Munich, Germany). The RNA was shipped on dry ice to Eurofins Genomics GmbH (Ebersberg, Germany). The subsequent analysis pipeline included rRNA depletion, library preparation (mRNA fragmentation, strand specific cDNA synthesis), Illumina paired end sequencing (2 x 150 bp, minimum 10 MB reads)), and bioinformatic analysis (mapping against the reference genome, identification and quantification of transcripts, pairwise comparison of expression levels and determination of significant fold differences) and was conducted by the company.

### Phylogenetic tree inference

For inference of a phylogenetic tree for sHdrR proteins and relatives, proteins were aligned using MAFFT (Katoh and Standley, 2013) and the alignment was manually trimmed. A maximum likelihood phylogeny was inferred iusing IQ-TREE v1.6.12 (Nguyen et al., 2015). Branch support was calculated by ultrafast bootstrap (2000 replicates) (Hoang et al., 2018). Finally, the tree was displayed using iTol (Letunic and Bork, 2021).

### Statistics and reproducibility

Experimental data are expressed as the mean ±standard deviation of the mean (SEM) of the number of tests stated for each experiment. All analysis was reproduced in at least three independent experiments. The significant difference between the two groups was analyzed using an independent student’s t-test; the p-value < 0.05 indicated statistical significance.

## RESULTS

### Distribution and phylogeny of sHdrR-related proteins

The *H. denitrificans* sHdrR (*Hd*sHdrR) protein is 125 amino acids in length, and a BlastP search identified *R. capsulatus* SqrR (Shimizu et al., 2017) as the most similar structurally characterized protein. *Hd*sHdrR is also similar to several other characterized transcriptional regulators, all of which belong to the same ArsR subfamily and include SoxR from *Pseudoaminobacter salicylatoxidans* (Mandal et al., 2007), *Paracoccus denitrificans* (Rother et al., 2005) and *H. denitrificans* (Li et al., 2023a), BigR from *Xylella fastidiosa* (Guimarães et al., 2011) and *Acinetobacter baumannii* (Walsh et al., 2020), YgaV from *E. coli* (Gueuné et al., 2008; Balasubramanian et al., 2022) and HlyU from *Vibrio cholerae* (Pis Diez et al., 2023) (Figure 1). All of these regulators share two conserved cysteine residues, Cys^50^ and Cys^116^ in the hyphomicrobial protein (Figure 1). The characterized proteins sense reactive sulfur species (RSS) and form an intraprotomer tetrasulfide bridge between the conserved cysteines when exposed to sulfane sulfur transpersulfidation donors (Giedroc et al., 2023; Li et al., 2023a).

**FIGURE 1.**
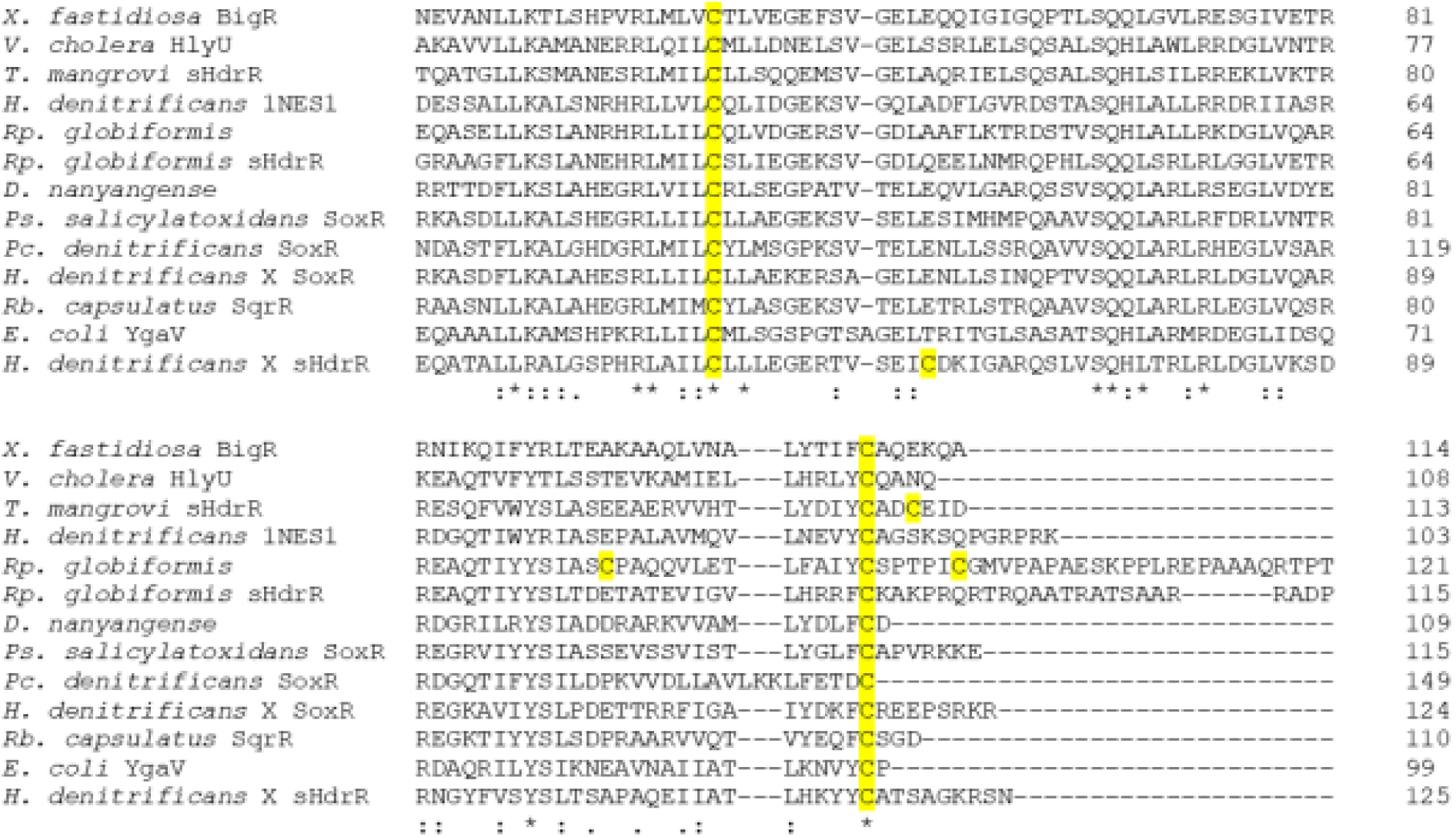
Amino sequence alignment of the central region of selected sHdrR homologs. Organism names, accession numbers/locus tags and references (if available) in the order of appearance: *Xylella fastidiosa* BigR, XF_0767 (Guimarães et al., 2011), *Vibrio cholera* HlyU, VC_A0642 (Mukherjee et al., 2014; Pis Diez et al., 2023); *Tsuneonella mangrovi*, CJO11_RS12710; *Hyphomicrobium denitrificans* 1NES1, HYPDE_25308 (Venkatramanan et al., 2013); *Rhodopila globiformis*, CCS01_RS26760 and *Rp. globiformis* sHdrR, CCS01_RS13140 (Imhoff et al., 2018); *Devosia nanyangense* HY834_20740 (He et al., 2021), *Pseudaminobacter salicylatoxidans* SoxR, WP_019171658 (Mandal et al., 2007); *Paracoccus denitrificans* SoxR, CAB94376 (Rother et al., 2005)*, H. denitrificans* X^T^ SoxR, Hden_0700 (Li et al., 2023a); *Rhodobacter capsulatus* SqrR, ADE85198 (Shimizu et al., 2017); *Escherichia coli* YgaV, b2667 (Gueuné et al., 2008); *H. denitrificans* X^T^ sHdrR, Hden_0682 (Li et al., 2023b; Li et al., 2024). An * (asterisk) indicates positions with identical residues. Cysteines are highlighted in yellow. Colons (:) and single dots (.) indicate conserved and semi-conserved amino acids, respectively.

In a previous publication, we performed a survey of the occurrence of genes for sHdrR-like proteins based on an automatically generated hidden Markov model (HMM) with the additional condition that the respective gene is located in the vicinity of a *shdr*-like gene cluster (Li et al., 2024). This revealed related transcriptional regulators in a wide variety of prokaryotes bearing the genetic potential for sulfur oxidation in the cytoplasm via the sHdr system. However, close examination of the amino acid sequences showed that many of the identified proteins do not contain the two conserved cysteine residues typical of sulfane-sulfur-responsive transcriptional repressors. This means not only that the HMM for sHdrR does not emphasize the conserved cysteines and must therefore be interpreted with caution, but also that the expression of *shdr* genes in many organisms involves ArsR-type regulators that rely on different modes of action.

In bacteria, transcriptional regulators often control the expression of genes located close to their own (Martinez-Antonio and Collado-Vides, 2003; Browning and Busby, 2004; Seshasayee et al., 2006). Therefore, we linked the proteins retrieved by a BlastP search using *Hd*sHdrR as a bait and the presence of the two conserved cysteines as a constraint with information about their genetic environment, and indeed we obtained revealing information (Supplementary Table 2, Figure 2). The general picture that emerged from our analysis is that sulfane sulfur-responsive ArsR-type regulators appear to act not only on the expression of genes for enzymes involved in the oxidation and/or detoxification of sulfur compounds, such as *sqr* for sulfide:quinone oxidoreductase, *sox* for thiosulfate oxidation in the periplasm, *pdo* for persulfide dioxygenase, *rhd* for rhodanese-type sulfur transferases or *shdr* for sulfite production in the cytoplasm, but also on target genes encoding membrane proteins that have been shown or are likely to be involved in the transport of sulfur compounds across the cytoplasmic membrane, such as efflux pumps of the resistance-nodulation-division (RND) family (Nikaido, 2011), sulfite exporters of the TauE family (Weinitschke et al., 2007), or SoxT and PmpAB, YeeE family transporters involved in the import of sulfur-containing ions (Gristwood et al., 2011; Tanaka et al., 2020; Ikei et al., 2024; Li et al., 2024) (Figure 2, Supplementary Table 2).

**FIGURE 2.**
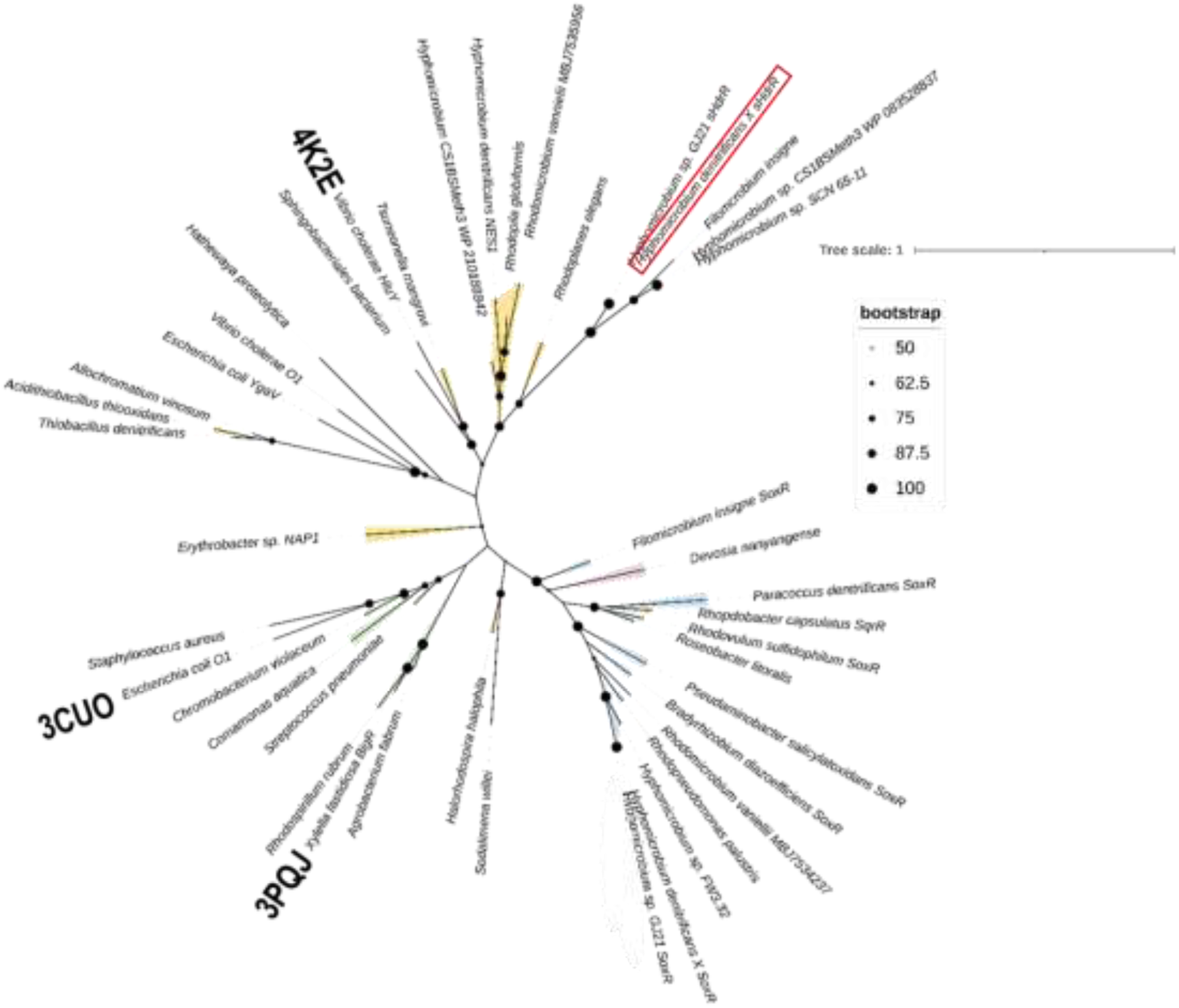
Unrooted phylogenetic tree for sulfane sulfur-responsive ArsR-type transcriptional regulators containing two conserved cysteine residues. Colored ranges indicate the occurrence of the respective gene in immediate vicinity of genes for resistance-nodulation-division (RND) transporters (beige), PmpAB-like YeeE family transporters (green), Sox proteins for thiosulfate oxidation (blue) or sHdr proteins for sulfur oxidation in the cytoplasm (purple). Note that in *Tsuneonella mangrovi* the *shdrR-*like gen is situated next to *shdr* genes and to genes for a RND transporter and that SoxR is encoded between sets of *sox* and RND-encoding genes in *Rhodopseuomonas palustris*. In both cases, only the vicinity to RND-encoding genes is indicated. The sHdrR protein from *H. denitrificans* X^T^ is highlighted by a red box. PDB codes are given for structurally characterized proteins. The tree was calculated with 2000 bootstrap resamplings using Ultrafast Bootstrap (Hoang et al., 2018) and IQ-Tree (Trifinopoulos et al., 2016; Minh et al., 2020). Bootstrap values between 50% and 100% are displayed as scaled circles at the branching points. Protein accession numbers, information on adjacent genes and references are available in Supplementary Table 2. The tree is available in Newick format as Supporting information (Supplementary data sHdrR and relatives tree.nwk).

Remarkably, *Hd*sHdr-related regulators with two conserved cysteines are rarely encoded in the vicinity of *shdr* genes and the few that fall into this category do not form a coherent phylogenetic clade (Supplementary Table 2, Figure 2). *Hd*sHdrR has a third cysteine, Cys^63^ (Figure 1). An equivalent of *Hd*sHdrR Cys^63^ is only present in a few cases (e.g. in the proteins from *Hyphomicrobium* sp. strains GJ21 and SCN 65-11). Neither is the absence of the third cysteine an indicator that the regulator is likely unable to act on *shdr* genes (due to its absence in the proteins from *Devosia nanyangense*, *Tsuneonella mangrovi* and *Rhodopila globiformis*, all of which are encoded in close proximity to *shdr* gene clusters), nor is its presence typical for ArsR-type proteins, which are likely to regulate *shdrR* genes, as it is present in the proteins of the cyanobacterium *Sodalimona willei* and the Gram-positive *Hathewaya proteolytica*. Both cannot oxidize sulfur compounds for dissimilatory purposes and do not contain *shdr* or *sox* genes (Figure 1, Supplementary Table 2).

### Importance of conserved sHdrR cysteines *in vivo*

In order to build a firm basis for the identification and characterization of sHdrR, we clarified whether the conserved cysteines (Cys^50^ and Cys^116^ in *Hd*sHdrR, Figure 1) are indeed necessary for proper function of the regulator *in vivo.* This was achieved by phenotypic characterization of *H. denitrificans* mutant strains that contained chromosomal replacements of the wild-type *shdrR* gene with variants encoding cysteine to serine exchanges of either one or both of the two conserved cysteines. All experiments reported in this and also previous studies (Li et al., 2023b; Li et al., 2023a) were performed using *H. denitrificans* Δ*tsdA* as the reference strain. This strain lacks the gene for thiosulfate dehydrogenase (TsdA). This enzyme catalyzes the oxidative formation of tetrathionate from two thiosulfate molecules. Thiosulfate dehydrogenase-deficient strains are unable to form tetrathionate and thus are perfectly suited for analyzing oxidation of the sulfur substrate to sulfite and eventually sulfate (Li et al., 2023b; Li et al., 2023a; Tanabe et al., 2023; Li et al., 2024; Tanabe et al., 2024). The *H. denitrificans* Δ*tsdA* reference strain excretes sulfite as an intermediate *en route* to sulfate, when methanol-grown cultures are provided with thiosulfate as an additional electron source. Sulfite is toxic and leads to growth retardation that is particularly impressive when cultures are inoculated with thiosulfate-induced cells ((Li et al., 2023b), also compare curves with filled circles in the upper panels of Figure 3). A Δ*tsdA* strain, that additionally lacks the transcriptional repressor *shdrR*, oxidizes thiosulfate without induction and accordingly, its growth rate slows down almost immediately when it is exposed to the sulfur compound ((Li et al., 2023b; Li et al., 2024), also compare curves with open boxes in the upper panels of Figure 3). When the *shdrR* gene is complemented *in cis*, the wild-type phenotype is reconstituted (Supplementary Figure 2).

**FIGURE 3.**
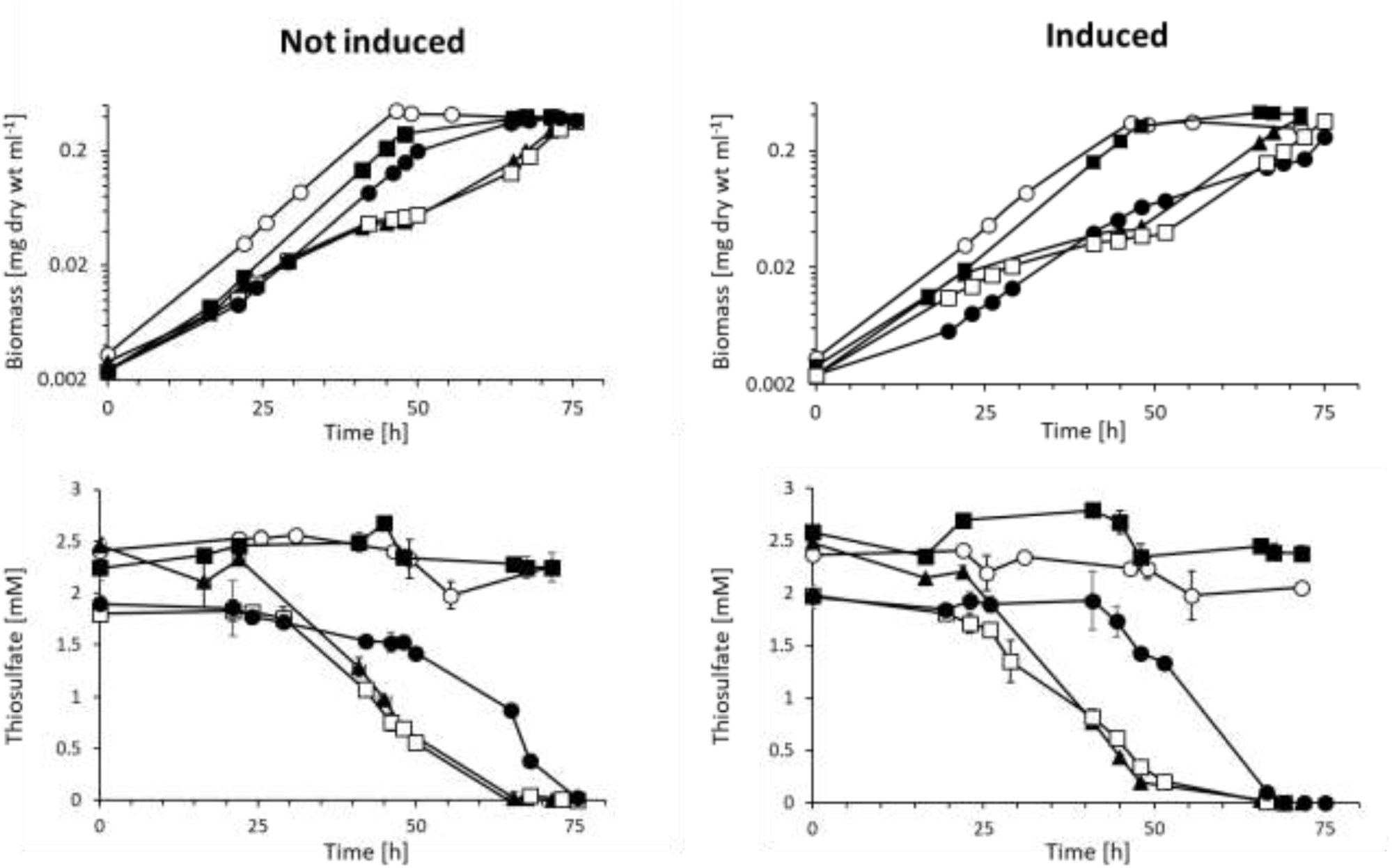
Growth and thiosulfate consumption of the *H. denitrificans* reference strain, a strain lacking sHdrR and strains producing sHdrR variants with cysteine to serine exchanges. *H. denitrificans* Δ*tsdA* (filled black circles) Δ*tsdA* Δ*shdrR* (open boxes), Δ*tsdA* sHdrR Cys^50^Ser filled boxes), Δ*tsdA* sHdrR Cys^116^Ser (filled triangles) and Δ*tsdA* sHdrR Cys^50^Ser Cys^116^Ser (open circles). All strains were grown in medium containing 24.4 mM methanol. In the “not induced” case, pre-cultures were grown without thiosulfate. In the “induced” case, pre cultures contained 2 mM thiosulfate. Thiosulfate concentrations for the different cultures are depicted in the lower panels. Symbol assignments are the same as in the upper panels. Error bars indicating SD are too small to be visible for determination of biomass. All studied strains grew equally well on methanol in the absence of thiosulfate (Supplementary Figure 1).

Like the *H. denitrificans* Δ*tsdA* Δ*shdrR* strain, the strain with sHdrR bearing a Cys^116^Ser exchange exhibited a high specific thiosulfate oxidation rate and a significantly reduced growth rate even without induction of pre-cultures (curves with filled triangles in Figure 3). The growth rate increased significantly after thiosulfate was consumed. This appears to mean that *in vivo* the sHdrR Cys^116^Ser variant protein has a decreased DNA binding and thus transcription repressing activity. In contrast, both, the mutant strain encoding the sHdrR Cys^50^Ser exchange and the mutant strain with both sHdrR cysteines replaced by serine, were unable to oxidize thiosulfate (curves with open circles and filled boxes in Figure 3). The most obvious conclusion here would be that these sHdrR repressor variants constitutively repress transcription, i.e. are always attached to their binding sites, and cannot react to the presence of oxidizable sulfur *in vivo*. However, we must take into account that the mutant strains studied still contain the fully functional SoxR regulatory protein, which we already know is the master regulator of sulfur oxidation in *H. denitrificans* (Li et al., 2024). Interactions with this protein *in vivo* may complicate conclusions about DNA binding capacity of the sHdrR variants. Irrespective of this, however, our experiments clearly demonstrate the relevance of the two conserved cysteines.

### Identification of genes controlled by sHdrR by RT-qPCR for different *H. denitrificans* strains

As a first step into the description of the sHdrR regulon, we performed RT-qPCR and determined the transcription levels of twelve signature genes in the *H. denitrificans* sulfur oxidation locus for the Δ*tsdA* Δ*shdrR* mutant in the absence of thiosulfate. The results were compared with previous results for the Δ*tsdA* reference strain in the absence and presence of thiosulfate, and with results for a Δ*tsdA* Δ*soxR* mutant in the absence of thiosulfate (Li et al., 2023a) (Figure 4a). In accordance with comparative mRNA-Seq analysis (Li et al., 2023a), our work with the reference strain had already shown a strong positive effect of thiosulfate on the expression of all genes residing in the respective region (Figure 4b), including two divergently transcribed *sox* operons for the periplasmic thiosulfate-oxidizing Sox system, the *soxT1A* operon which encodes a sulfane sulfur import machinery, the *dsrE3C-shdr-lbpA* genes responsible for cytoplasmic sulfane sulfur transfer and its oxidation to sulfite and the *lip* genes for assembly of lipoate on the LbpA proteins. The only unaffected genes were those for the transcriptional repressors, sHdrR and SoxR, and the transporter SoxT1B, that acts as a signal transduction unit for SoxR (Li et al., 2024). For the hyphomicrobial sulfur oxidation signature genes, the absence of SoxR had similar effects as the presence of thiosulfate, resulting in, but not limited to, high constitutive transcription rates for the *sox* and *shdr* genes (Figure 4a). As observed for SoxR, the absence of sHdrR led to constitutive transcription of genes relevant for oxidative sulfur metabolism. However, our qRT-PCR analyses showed that sHdrR affects only a subset of the genes affected by SoxR and that this sHdrR-controlled subset does not include the genes in the divergently transcribed *soxYZ* and *soxAX* loci (Figure 4a).

**FIGURE 4.**
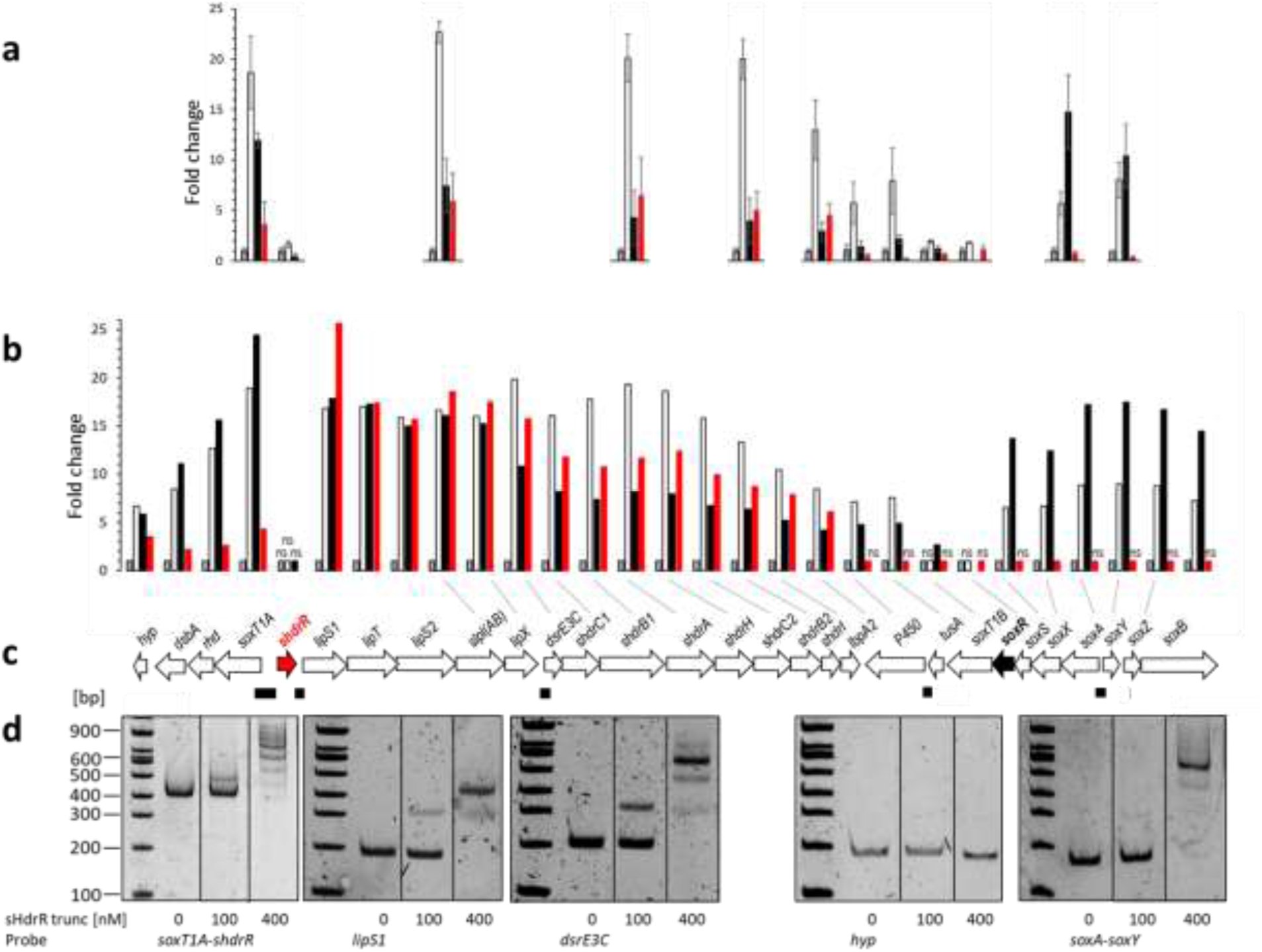
Regulation of the *shdr-sox* region in *H. denitrificans*. **(a)**. Relative mRNA levels of twelve genes located in the *shdr-sox* region (depicted in panel **(c)**) from *H. denitrificans* for the Δ*tsdA* reference strain in the absence (gray columns) and presence of thiosulfate (white columns), as assessed by RT-qPCR. Results for *H. denitrificans* Δ*tsdA* Δ*soxR* and *H. denitrificans* Δ*tsdA* Δ*shdrR* are shown by black and red columns, respectively. Results were adjusted using *H. denitrifcans rpoB*, which encodes the β-subunit of RNA polymerase, as an endogenous reference according to (Martineau et al., 2015). All cultures were harvested in the exponential growth phase. Three parallel experiments were performed to obtain averages and standard deviation. **(b)** Transcript abundance changes of genes encoding enzymes involved in thiosulfate oxidation in the same strains as in **(a)** as assessed by mRNA-Seq analysis. The same color coding applies. The experiments were conducted in duplicate, each time using mRNA preparations from two different cultures. Adjusted *p* values for statistically significant changes were all below 0.001 (Supplementary Table 3). ns, not significant. **(c)** DNA regions tested in EMSA assays for sHdrR binding are indicated as black rectangles below the hyphomicrobial *shdr-sox* genetic island. Fragment sizes: 362 bp for the *soxT1A-shdrR* intergenic region, 177 bp and 173 bp for the regions upstream of *lipS1* and *dsrE3C*, respectively. The fragments downstream of *tusA* and between *soxA* and *soxY* had sizes of 176 bp and 151 bp, respectively. **(d)** EMSA analysis of Strep-tagged sHdr-trunc with upstream promoter sequence probes of sulfur oxidation related genes as specified in **(c)**. 17 nM DNA probes were incubated with different amounts of sHdr-trunc (100 and 400 nM). Vertical lines separate samples that were run on the same gel but were not directly adjacent.

The finding that sHdrR does not exert a significant effect on *sox* gene expression *in vivo*, prompted us to analyse probable DNA binding regions. To that end, we inspected the same intergenic regions within the hyphomicrobial sulfur oxidation locus that had already been tested as probes for SoxR (Li et al., 2023b) and used them in electrophoretic mobility shift assays (EMSA) with sHdrR (Figure 4c,d). These experiments were performed with a carboxy-terminally His-tagged and amino-terminally truncated version of sHdrR produced in *E. coli*, that lacked the first 25 amino acids of the wild-type protein. The truncation was necessary because the full-length sHdrR proved to be very unstable, as documented by mass spectrometry, whereas the truncated variant could be used for a period of seven days when kept on ice. The truncated sHdrR variant fully encompasses the central conserved region shown in Figure 1 and binds effectively to all but the control DNA probe (Figure 4d). The latter is a 176-bp fragment located upstream of Hden_0697, encoding a putative cytochrome P450 that, unlike the other fragments, does not contain prominent inverted or direct repeats (Figure 4c). All four of the DNA fragments that were bound by sHdrR had previously been shown to contain binding sites for SoxR (Li et al., 2023b). Surprisingly, the intergenic region between the divergently oriented *soxA* and *soxY* genes is recognized by sHdrR *in vitro* (Figure 4d), although a lack of sHdrR has no significant effect on the transcription of these genes (Figure 4a). Together, our observations point at an intimate interplay of the two related repressor proteins *in vivo* that might be governed by different binding affinities and/or include heterocomplex formation.

### Global analysis of the sHdrR and SoxR regulons by RNA-seq analysis of different *H. denitrificans* strains

In the next step, we sought to identify the regulons of the intertwined transcriptional repressors SoxR and sHdrR on a global basis and extended previous genome-wide mRNA-Seq data for the Δ*tsdA* reference strain to *H. denitrificans* strains lacking either the genes for sHdrR or SoxR. We compared the mRNA abundance in these two strains to the mRNA abundance in the reference strain in the absence of thiosulfate. In addition, the data set for the reference strain in the presence of thiosulfate (Li et al., 2024) was integrated in the analyses. The number of identified mRNAs was virtually identical in all cases and ranged between 95 to 96% of the 3529 predicted genes.

Volcano plots (Supplementary Figure 3) offer a comparative visual assessment of total gene expression patterns for each mutant relative to the wild type in the absence and in the presence of thiosulfate. The lack of the repressors sHdrR and SoxR affected the abundance of a total of 165 (4.8%) and 170 (5.0%) of the detected mRNAs, respectively, while the presence of thiosulfate affected the transcription of 136 (4.1%) genes in the reference strain. The genes affected by the lack of the two repressors exhibit a high overlap of 138 genes (83.6% and 81.2% of all genes affected by sHdrR and SoxR, respectively) (Figure 5a). Furthermore, 85 genes (65%) that are affected by the lack of either one or both of the repressors overlap with the genes affected by thiosulfate in the reference strain. We can therefore confidently state that SoxR and sHdrR are intimately involved in the cells’ response to the availability of the oxidizable sulfur substrate thiosulfate.

**FIGURE 5.**
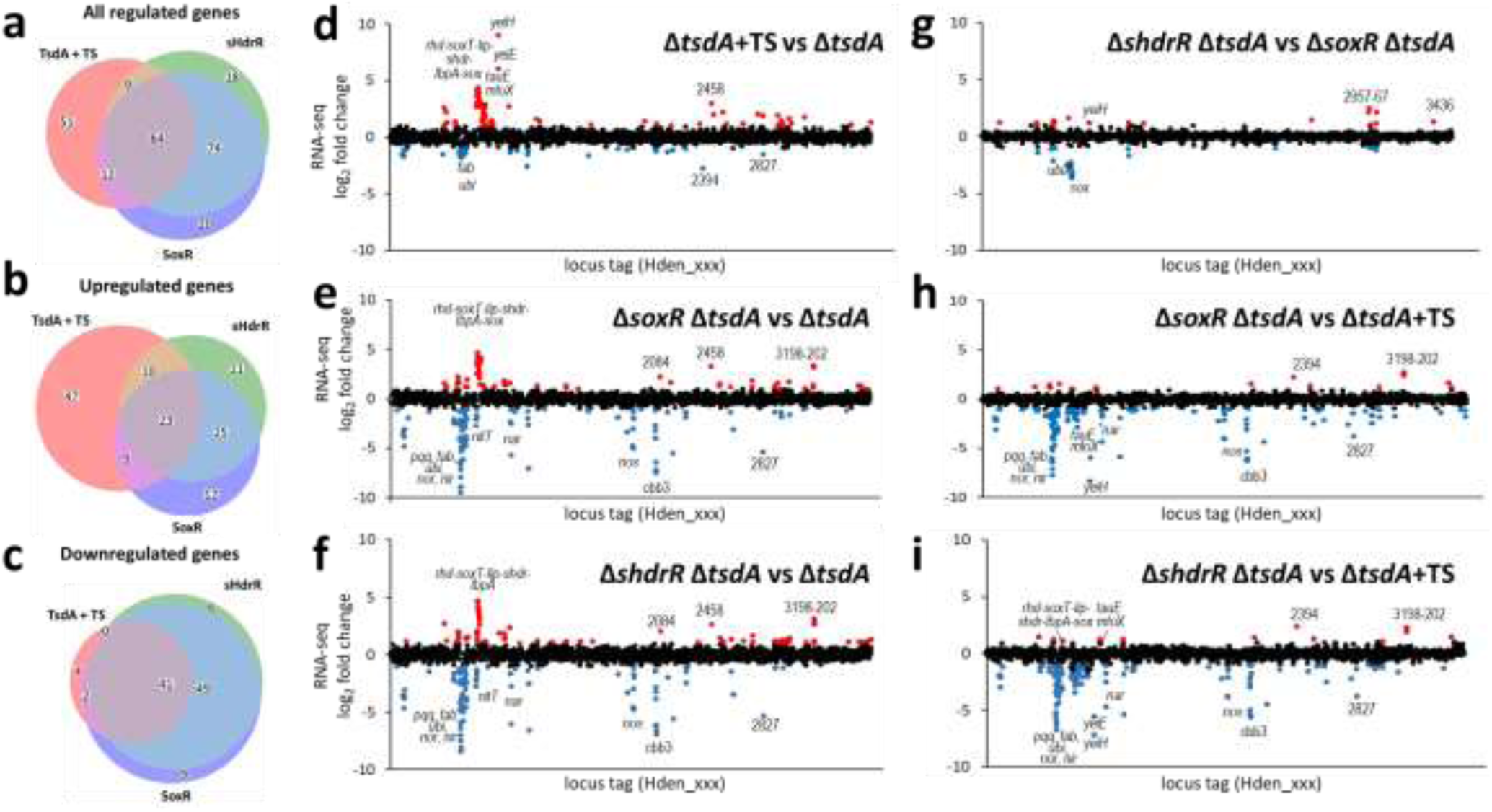
Global analysis of the sHdrR and SoxR regulons in *H denitrificans* by RNA-seq analysis. **(a to c**) Scaled Venn diagrams displaying the number differentially expressed genes from mRNA-Seq analysis of *H. denitrificans* strain Δ*tsdA* in the presence of thiosulfate (TsdA + TS, red), and strains Δ*tsdA* Δ*soxR* (SoxR, blue) and Δ*tsdA* Δ*shdrR* (sHdrR, green) in the absence of thiosulfate compared to *H. denitrificans* strain Δ*tsdA* in the absence of thiosulfate. The number of differentially expressed genes in each group is represented by the size of each circle, and overlapping areas indicate genes shared between the strains/conditions. Genes meeting a log2-fold change of >2 and an adjusted p-value of <0.001 were considered differentially expressed. Numbers in each area are given for all regulated genes **(a),** all upregulated genes **(b)** and all downrgulated genes **(c)**. **(d to i)** Fold change in transcription for *H. denitrificans* Δ*tsdA* + thiosulfate (TS) versus Δ*tsdA* untreated **(d)**. Δ*tsdA* Δ*soxR* untreated versus Δ*tsdA* untreated **(e)**. Δ*tsdA* Δ*shdrR* untreated versus Δ*tsdA* untreated **(f)**. Δ*shdrR* Δ*tsdA* versus Δ*tsdA* Δ*soxR* **(g)**. Δ*tsdA* Δ*soxR* untreated versus Δ*tsdA* treated **(h)**. Δ*shdrR* Δ*tsdA* untreated versus Δ*tsdA* treated **(i)**. Genes with a significant fold change (*p*<0.001) in two biological replicates are highlighted (red for a >2-fold increase or blue for a >2-fold decrease.

Based on current models of how ArsR-type repressors control gene expression (Busenlehner et al., 2003; Ren et al., 2017; Roy et al., 2018), removing the repressor proteins (i.e. deletion mutations as in the current study) should result in constitutive expression of genes that are normally repressed by that protein in the absence of the de-repressing effector molecule. Decreased expression would indicate that the gene is activated (directly or indirectly) by the respective transcriptional regulator. The presence of thiosulfate positively influences (fold change >2) the transcription of 89 genes in the reference strain, while a lower number of genes (47) is negatively affected (Li et al., 2024). Both positive and negative changes are also observed for the SoxR and the sHdrR-deficient mutant strains (Figure 5b,c). In the *H. denitrificans* Δ*tsdA* Δ*soxR* and Δ*tsdA* Δ*shdrR* strains, 69 genes are significantly overexpressed respectively, with an overlap of 69.6% of the genes (Figure 5b). The overlap with transcripts of higher abundance in the presence of thiosulfate in the reference strain is 37.1% and 36.0%, respectively (Figure 5b,d, Supplementary Table 3). Conspicuously, there is a much higher overlap between the transcription of genes negatively affected by thiosulfate or the absence of either one of the repressors than seen for the genes with transcription increases (compare Figures 5b and 5c). While for the upregulated cases, 47 (52.8%) are induced by thiosulfate in the reference strain but do not show consequences upon deletion of the repressor genes, this is observed for only 4 (8.5%) of the downregulated cases. In addition, there are very few genes for which transcriptional downregulation is caused by the absence of only one of the regulators (Figure 5c). Only in 6.7% and 12.2% of cases, respectively, is the effect exclusively due to the absence of sHdrR or SoxR, while for genes with increased transcription the proportion amounts to 30.4% in both cases.

Here, we focus first on the genes whose transcription is strongly increased in the absence of one or both of the two transcriptional regulators as well as in the presence of thio-sulfate in the reference strain (Figure 5d-f). Most of these genes encode enzymes involved in oxidative sulfur metabolism (Supplementary Table 3, Figure 6a). In full agreement with the RT-qPCR analysis (Figure 4a), the transcription of the genes of the *soxT1A* operon, the *dsrE3C-shdr-lbpA* and the *lip* genes, which together encode proteins for sulfane sulfur import and its oxidation to sulfite, was strongly affected by the absence of either one, the sHdrR or the SoxR repressor (Figures 4b, 5e,f). The situation was completely different for the *sox-tusA-p450* genes. Here, sHdrR deficiency had only a marginal effect, if any, which is again fully consistent with our RT-qPCR results (Figures 4a,b, 5f). The genes for the transcriptional repressors themselves and that for the signal transducing SoxT1B membrane protein were barely affected by thiosulfate or the absence of the other repressor. There is a further conspicuous gene, Hden_2458, the transcription of which is increased in the presence of thiosulfate as well as in the absence of both repressor proteins (Figure 5d-f). It encodes a small, soluble, cytoplasmic hypothetical protein of 46 amino acids with no known function.

**Figure 6.**
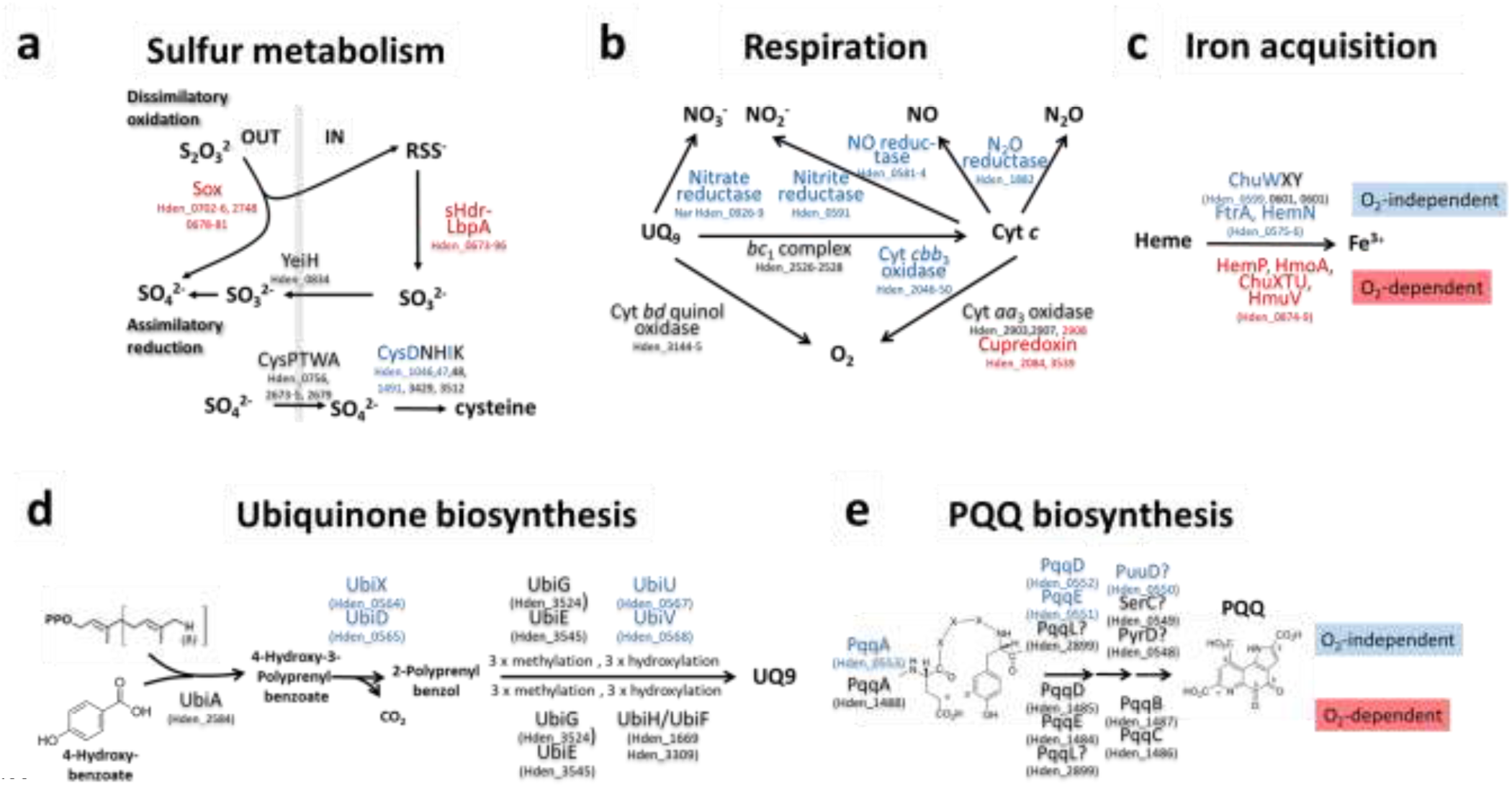
Effects of the absence of sHdrR or SoxR on central metabolic pathways in *H. denitrificans* X^T^. Proteins whose genes showed increased (>2fold) transcript abundance either in *H. denitrificans* strain Δ*tsdA* Δ*shdrR* or in strain Δ*tsdA* Δ*soxR* or in both when compared to the *H. denitrificans* Δ*tsdA* reference strain in the absence of thiosulfate are printed in red. Proteins whose genes showed decreased (<0.5fold) transcript abundance in either of the three strains are printed in blue. Black letters indicate that significant changes in transcript abundance were not observed. **(a)** Dissimilatory oxidative and assimilatory reductive sulfur metabolism. **(b)** Electron delivery pathways to respiratory electron acceptors **(c)** Iron acquisition by heme degradation and iron release **(d)** Oxygen-dependent and oxygen independent biquinone biosynthesis. In *E. coli,* ubiquinone biosynthesis starts from 4-hydroxybenzoate, that is produced from chorismate (Abby et al., 2020), a step catalyzed by chorismate-pyruvate lyase (UbiC) (Pelosi et al., 2016). We are currently unable to explain how 4-hydroxybenzoic acid is synthesized in *H. denitrificans* and whether it is a mandatory precursor at all. *H. denitrificans* does not encode either a UbiC homolog or a homolog of XanB2, an unrelated chorismatase that fills the role of UbiC in *Xanthomonas campestris* (Zhou et al., 2013). In addition, it has been reported that alphaproteobacterial UbiA can accept *p*-amino-benzoic acid as a substrate for prenylation (Xie et al., 2015; Degli Esposti, 2017). **(e)** (Proposed) pathways of PQQ synthesis in the absence or presence of oxygen.

When we look at the genes with increased transcript abundance only in the regulator-deficient mutants, several sets of genes stand out that form a functionally coherent group (Figure 5d-f, Supplementary Table 3, Figure 6c). The encoded enzymes all play a role in iron acquisition via uptake and subsequent degradation of hemin (Wandersman and Delepelaire, 2004) and include Hden_0541 and Hden_0875, two putative oxygen-dependent heme-degrading HmoA monoxygenases (Frankenberg-Dinkel, 2004). Key subunits of the sulfur-oxidizing Sox and sHdr systems contain either heme or iron-sulfur sulfur clusters, and it is therefore not surprising that cells prepare for thiosulfate oxidation by taking steps to ensure that these enzymes are equipped with the necessary prosthetic groups.

Other conspicuous positive changes in abundance seen only in the repressor-deficient mutants include genes for proteins involved in electron transport/aerobic respiration (Supplementary Table 3, Figure 6b). Among these are Hden_2084 and Hden_3539, both of which encode periplasmic pseudoazurins/cupredoxins. While the small copper-binding cupredoxins are best known as electron donors to the denitrification pathway (Kataoka et al., 2004; Impagliazzo et al., 2005; Fujita et al., 2012), members of the protein family can also be required for cytochrome *c* oxidase respiratory function under aerobic conditions (Castelle et al., 2010).

The two genes that show the highest abundance changes (526- and 66-fold, respectively) in the reference strain upon addition of thiosulfate, encode a sulfite exporter (Hden_0834, YeiH) and a LysR family transcriptional regulator (Hden_0835) (Li et al., 2024). Surprisingly, the transcription of these two genes is not affected by the removal of sHdrR or SoxR. Closer inspection of the LysR-type protein revealed that its most closely related structurally and biochemically characterized homologs are NdhR (or CcmR) from *Synechocystis* PCC6803 (5Y2V (Jiang et al., 2018) and YeiE from *Cronobacter sakazaki* (7ERQ_A (Hong et al., 2022)). While 2-phosphoglycolate is an inducer for NdhR, YeiE serves as a global virulence regulator in *C. sakazakii*, binds sulfite as an effector and has a central role in defending against sulfite toxicity (Hong et al., 2022). The Hden_0835 derived protein shares five of seven sulfite-interacting residues with *C. sakazakii* YeiH. Notably, these include Glu^145^, which is responsible for discriminating between sulfite and sulfate (Supplementary Figure 4). In contrast, of the seven residues in NdhR that interact with 2-phosphoglycolate, only three are present in the *H. denitrificans* protein. Toxic sulfite is formed as an intermediate during thiosulfate oxidation by *H. denitrificans* (Li et al., 2023b) and is effectively excreted into the medium, probably mainly by the action of the YeiH exporter (Li et al., 2024). It is likely that the YeiE protein derived from Hden_0835-recognizes sulfite and activates transcription of the neighboring *yeiH* gene upon binding of the inducer. This is only seen in the *H. denitrificans* Δ*tsdA* reference strain because the sHdrR- and SoxR-deficient strains were cultivated in the absence of thiosulfate. There is another set of genes, Hden_0719 to Hden_0748, for which drastic changes in abundance occur when the reference strain is exposed to thiosulfate, but whose transcription is not triggered by the absence of sHdrR or SoxR, i.e. these genes do not belong to the regulons of these two repressors. The affected genes include those for a putative dimethylsulfide (DMS) monooxygenase and for methanethiol oxidase, MtoX (Eyice et al., 2017). In *H. denitrificans* thiosulfate is formed during DMS degradation, which occurs with methanethiol as an intermediate product (Koch and Dahl, 2018). We conclude that cells that sense thiosulfate are stimulated to prepare for degradation of the environmentally abundant organosulfur compound.

We have previously reported that transcription of a number of genes decreases when the *H. denitrificans* Δ*tsdA* reference strain is exposed to thiosulfate and that the encoded proteins include those of assimilatory sulfate reduction, since thiosulfate can serve as a source of reduced sulfur (Li et al., 2024). A lower transcript abundance for the genes for sulfate adenylyltransferase CysDN, which catalyzes the activation of sulfate to adenosine-5’-phosphosulfate, is also observed in the repressor-deficient *H. denitrificans* strains (Supplemenmtary Table 4, Figure 6a), but the response to the lack of the repressors extends far beyond sulfur assimilation to central energy metabolism (Figure 5d-I, Figure 6a-e).

First, transcript abundance of the genes for cytochrome *c* oxidase of type *cbb*_3_ is greatly reduced (Supplementary Table4, Figure 6b). This oxidase is adapted to low oxygen concentrations and plays a crucial role in microaerobic respiration (de Gier et al., 1996; Pitcher and Watmough, 2004). It has a higher affinity to oxygen and is less efficient in proton pumping compared to the *aa*_3_-type cytochrome *c* oxidase, the transcription of which remains unaffected. Second, transcript abundance for virtually all genes underlying nitrate respiration is drastically reduced when either sHdr or SoxR are absent. Affected units include nitrate and nitrite transporters/antiporters (NitT, Nark), nitrate reductase (Nar), nitrite reductase (Nir), nitric oxide reductase (Nor), nitrous oxide reductase (Nos), electron carriers involved in denitrification and redox balance (cytochromes *c*, cupredoxin, NosR/NirI) as well as nitrate/nitrite and NO responsive regulators (NarQL, NnrS) (Figure 5e,f, Supplementary Table 4, Figure 6b). It should be noted that the same differences are seen when the regulator-deficient mutant strains are compared with the reference strain in the presence ot thiosulfate (Figure 5h,i).

The diminished transcript abundance of genes for enzymes involved in ubiquinone (UQ) biosynthesis (Hden_0564 to 0568, UbiXDTUV) is directly related to denitrification, since ubiquinol serves as an electron donor for nitrate reduction. The first step in ubiquinone biosynthesis is prenylation of 4-hydroxy benzoic acid by the membrane-bound enzyme UbiA (Hden_2584), followed by decarboxylation (catalyzed by the UbiX-UbiD system), three hydroxylations, two *O*-methylations (catalyzed by UbiG) and one *C*-methylation (catalyzed by UbiE) (Figure 6d) (Aussel et al., 2014). Under anaerobic conditions, the hydroxylation reactions are achieved by the oxygen-independent UbiT, UbiV and UbiU proteins (Pelosi et al., 2019), all of which are essential for denitrification in *Pseudomonas aeruginosa* (Vo et al., 2020). It thus appears that oxygen-independent UQ synthesis is co-regulated with denitrification in *H. denitrificans*. It is important to note that the organism contains only UQ_9_, an ubiquinone with a side-chain containing nine prenyl residues (Urakami and Komagata, 1979, 1987). To the best of our knowledge, menaquinone has not been detected in any *Hyphomicrobium* species. Accordingly, neither homologs of the *E. coli* menaquinone biosynthetic pathway nor enzymes of the alternative so-called futasoline pathway (Dairi, 2012) are encoded in *H. denitrificans*. Since UQ_9_ is the only quinone, it must also be made available for aerobic respiration. In fact, two UbiH/UibF-like monooxygenases (Hden_1669, Hden_3309) are suitable for the catalysis of the hydroxylation reactions on the aromatic ring in the presence of oxygen. Other proteobacteria also contain only two or even only one of these enzymes with relatively broad regioselectivity instead of the three specifically acting prototypical *E. coli* enyzmes (Pelosi et al., 2016). *H. denitrificans* has only one copy each of *ubiD* and *ubiX* and these are hardly transcribed in our sHdrR- and SoxR deficient mutant strains even when grown under full oxygen tension (Supplementary Table 4, Figure 6d), suggesting that UbiD and UbiX may be replaced by so far unidentified non-orthologous proteins under aerobic conditions. The existence of alternative decarboxylases is supported by the fact that some ubiquinone-containing Alphaproteobacteria lack *ubiD* and *ubiX* completely. In addition, the decarboxylating enzyme has not yet been identified in mitochondria (Guerra and Pagliarini, 2023).

The clustered genes for oxygen-independent UQ_9_ biosynthesis (Hden_0564-571) and nitric oxide reductase plus accessory and regulatory proteins (Hden_0572-0595) are preceded by two other gene sets for which transcript abundance decreases significantly in the absence of either SoxR or sHdrR. The affected genes appear to be part of two divergently transcribed operons, Hden_0561-0563 and Hden_ 0560-0546, with the latter encoding a NnrS family protein (Gaimster et al., 2018). NnrS was initially described in *Rhodobacter* as a heme- and copper containing transmembrane protein (Kwiatkowski et al., 1997) and contributes to nitrosative stress tolerance (Stern et al., 2013). The proteins encoded by Hden_0554 to Hden_0563 all appear to be related to fatty acid biosynthesis and include four genes for 3-oxoacyl-ACP-[acylcarrier protein]-synthase II or possibly 3-oxoacyl-ACPsynthase I, FabF, one each for FabG (3-oxoacyl-ACP reductase) and FabZ (beta-hydroxyacyl-ACP dehydratase) and two for acyl carrier proteins (Hden_0560, 0563). For all of these genes, *H. denitrificans* has at least one additional copy residing somewhere else on the genome. The last step of the elongation cycle during fatty acid synthesis is catalyzed by enoyl-ACP reductase (FabI). However, the canonical hyphomicrobial FabI enzyme, Hden_1970, which is a member of the short-chain dehydrogenase/reductase superfamily member, does not have a counterpart in the gene cluster underlying SoxR/sHdrR control. We speculate that the product of Hden_0556, annotated as alcohol dehydrogenase, performs this function under anaerobic/denitrifying conditions. It is well known that some bacterial species contain additional enoyl-ACP reductases (Massengo-Tiasse and Cronan, 2009; Hopf et al., 2022). In summary, we suggest that the mentioned gene products work together in a fatty acid biosynthesis pathway that is especially efficient in/designed for anaerobic/denitrifying conditions.

Hden_0551-0553 encode proteins dedicated to biosynthesis of pyrroloquinoline quinone (PQQ). PQQ is a cofactor of periplasmic quinoprotein dehydrogenases such as cytochrome *c-*dependent methanol dehydrogenase, an enzyme of major importance to *H. denitrificans* when it grows on methanol (Duine and Frank, 1980b, a; Dijkstra et al., 1989). PQQ is synthesized from a precursor peptide, PqqA, with the conserved sequence motif E-X_3_-Y (Cordell and Daley, 2022) (Figure 6e). PqqA is bound by PqqD (Latham et al., 2015), and bond formation between the glutamate and tyrosine C9 atoms is catalyzed by PqqE. The next step involves the cleavage of the structure from PqqA, which is catalyzed by PqqF/PqqG /PqqH and/or other proteases (Cordell and Daley, 2022). PqqB probably hydroxylates and oxidizes the Glu-Tyr dipeptide yielding 3a-(2-amino-2-carboxyethyl)-4,5-dioxo-4,5,6,7,8,9-hexahydroquinoline-7,9-di-carboxylic acid (AHQQ). The last step is ring cyclization and eight-electron oxidation catalyzed by PqqC. The reaction includes four oxidative steps requiring molecular O_2_ and hydrogen peroxide (H_2_O_2_) (Bonnot et al., 2013). *H. denitrificans* X^T^ contains the full gene set for PQQ synthesis under aerobic conditions (PqqABCDE, Hden_1484-1488). Genes for PqqF/G/H are not present, but their function could be taken over by PqqL (Hden_2899) or other peptidases (Grinter et al., 2019; Cordell and Daley, 2022). Second copies for *pqqA*, *pqqD*, and *pqqE* are located in the gene cluster that is under SoxR/sHdrR control (Hden_0552 to 0553), and it is tempting to speculate that the final steps of PQQ biosynthesis in the absence of oxygen may be encoded in the same transcriptional unit and undergo the same transcriptional regulation. There is some circumstantial support for this suggestion: The products of Hden_0550 and Hden_0548 are similar to the N-terminal domain with four transmembrane helices of cytochrome *c* urate oxidase, PuuD (Doniselli et al., 2015), and to dihydroorotate oxidase, PyrD (Larsen and Jensen, 1985), respectively, both of which catalyze reactions on carbon- and nitrogen-containing heterocycles with certain structural similarities to precursors of PQQ. Hden_0549 encodes a putative phosphoserine aminotransferase, SerC (Duncan and Coggins, 1986). Clusters of a *pqqAED-puuD-serC-pyrD* are not only present in addition to classical *pqqABCDE* clusters in other *Hyphomicrobium* species, i.e. *H. nitrativorans*, but also in the Gammaproteobacteria *Halomonas sulfidovorans* strain MCCC 1A13718 (locus tags for the two *pqqE* genes: HNO53_16555 and HNO53_16620), *Stutzerimonas* (former *Pseudomonas*) *stutzeri* (*pqqE* genes in strain CGMCC 22915: NPN27_09805 and NPN27_22915) and *Methylophaga nitratireducenticrescens* JAM1 (*pqqE* genes: Q7A_453 and Q7A_868), and the Betaproteobacterium *Azoarcus* sp. DN11 (*pqqE* genes: CDA09_04515 and CDA09_08755). All of these organisms are capable of a denitrifying metabolism (Kasai et al., 2007; García-Valdés et al., 2010; Martineau et al., 2013; Wang and Shao, 2021) and may therefore be prepared for PQQ synthesis under these conditions. *Klebsiella pneumonia* and *Pseudomonas aeruginosa* are counterexamples. They contain only the canonical *pqqABCDEF/H* genes, and PQQ is not synthesized during anaerobic growth, although the *pqq* gene set is transcribed (Velterop et al., 1995; Gliese et al., 2010).

The last set of genes that deserves attention due to strong decreases in transcript abundance in the sHdrR and SoxR-deficient mutant strains, are those encoding proteins needed for heme degradation and iron acquisition under anaerobic conditions. FtrA (Hden_0575) is a periplasmic iron protein, HemN (Hden_0576) serves as an oxygen-independent coproporphyrinogen III oxidase (Layer et al., 2002) and ChuW (Hden_0599) is a radical S-adenosylmethionine methyltransferase that catalyzes a radical-mediated mechanism facilitating iron liberation and the production of a tetrapyrrole product called “anaerobilin” (LaMattina et al., 2016). It serves as a substrate for ChuY, an anaerobilin reductase (Hden_0601), possibly also involving ChuX, a putative heme binding protein (Hden_0600). The transcript abundance for these two genes is not affected in the repressor-negative mutants.

## DISCUSSION

Here, we collected information on sHdrR, an ArsR-type regulator that functions as a transcriptional repressor of genes encoding enzymes involved in the oxidation of thiosulfate as a supplemental electron donor in *H. denitrificans*. Hyphomicrobial sHdrR belongs to a family of sulfane-sensitive regulators characterized by two conserved and essential cysteines, which also includes SoxR from the same organism. SoxR and sHdrR are homologous proteins. DNA-binding studies and expression analyses using RT-qPCR and RNA-Seq techniques show that both are directly involved in the control of sulfur oxidation. While both proteins bind to the same DNA fragments upstream of sulfur oxidation-related genes *in vitro*, the removal of each individual regulator has overlapping but non-identical effects on the transcription of these genes *in vivo*. sHdrR regulates only a subset of SoxR-dependent genes, and this subset does not include the *sox* genes. These encode the enzymes that catalyze the initial steps of thiosulfate oxidation in the periplasm (Fig. 6a). Rather, in conjunction with SoxR, sHdrR is responsible for releasing transcriptional repression of genes for enzymes further downstream in thiosulfate degradation, i.e., import into the cytoplasm and oxidation to sulfite in this compartment (Fig. 6a). Given the close sequence similarity of SoxR and sHdrR, it is likely that their mechanism of action is similar, which would imply that the formation of a bridge of one to three sulfur atoms between the sulfur atoms of two conserved cysteine residues leads to a conformational change and unbinding of DNA (Li et al., 2023a). Whether and how the two transcriptional repressors compete for their binding sites or whether they even form heterocomplexes, cannot be answered on the basis of the available data. One possible model would be that SoxR and sHdrR co-repress their target promoters by binding as two separate homodimers. An alternative model would be that these two proteins repress their target promoters as a functional SoxR-sHdrR heterodimer. Heterodimerization of transcription factors is prevalent as a regulatory mechanism in eukaryotes (Remenyi et al., 2004) but is rare in bacteria. It has for example been reported for the LuxR-type transcription factor RcsB that forms heterodimers with several different auxiliary proteins. Like RcsB all of these carry a FixJ/NarL-type helix-turn-helix DNA binding motif (Kelm et al., 1997; Wehland and Bernhard, 2000; Venkatesh et al., 2010; Pannen et al., 2016). Another example is the interaction of BldM and WhiI from *Streptomyces*. Here, a BldM homodimer activates transcription of genes for early stages of development, while a BldM-WhiI heterodimer activates genes required for later stages (Al-Bassam et al., 2014).

Our global RNA-seq analyses of the *H. denitrificans* Δ*tsdA* reference strain in comparison with sHdrR- and SoxR-deficient strains have yielded detailed insights into the regulons of these sulfur-responsive transcriptional regulators. These regulons encompass not only genes for sulfur metabolism, but go considerably further than expected. This includes a drastic increase in the transcription of genes whose products are responsible for the availability of iron under aerobic conditions in the absence of one of the two repressor proteins. This finding underscores the critical need for a collaborative action of both proteins to maintain the transcription of the relevant genes at the optimal level. Comparable increases in abundance are not triggered by thiosulfate in the reference strain if it grows on the same defined medium with sufficient trace elements. It is thus evident that mutants lacking a single repressor are incapable of responding to iron availability. A logical explanation for this phenomenon is that the two repressors, functioning in conjunction, control the production of an iron-responsive regulator.

The most surprising result of our study is the unexpectedly large group of genes whose transcription is reduced by factors of up to 1000 in the absence of either sHdrR or SoxR (Figs. 5 and 6, Supplementary Table 4). While a comprehensive mechanistic explanation remains elusive, it is evident that the presence of both repressor proteins is imperative to prevent the near-total cessation of transcription of these genes. It is irrelevant which of the two repressor proteins is missing. In this context, we therefore observe a seemingly paradoxical phenomenon: repressor proteins are necessary for maintaining normal transcription levels. This effect is not observed in the *H. denitrificans ΔtsdA* reference strain when growing on thiosulfate, i.e., under conditions where neither of the two regulators can bind to its DNA target structures. On this basis, we once again conclude that cooperation between the two repressor proteins is indispensable for proper function *in vivo*. Furthermore, we suggest that, as with the regulation of iron availability, this is probably not a direct effect but rather an indirect one exerted via the control of other transcription factors. One explanation would be that the simultaneous presence of both repressor proteins is necessary to maintain the suppression of a stronger repressor. In the simultaneous presence of SoxR and sHdrR, the transcription of target genes would be sustained at a normal level due to the absence of the stronger repressor. However, if the repression of the controlled repressor is compromised due to the absence of either SoxR or sHdrR, then larger amounts of the stronger repressor protein can be formed, potentially leading to significant repression of the transcription of target genes. Alternative explanations include the indirect joint influence of SoxR and sHdrR on the transcription of an activator protein or participation in a feedback loop within a larger regulatory network.

The products of virtually all genes whose transcription is negatively affected in the absence of SoxR or sHdrR share a common overarching feature: their involvement in anaerobic metabolic pathways, particularly energy conservation in the absence of oxygen. It has long been known that *H. denitrificans* can grow on methanol as a carbon and electron source with respiration on nitrate as an electron acceptor. During aerobic growth of our regulator-deficient mutants, we observe a negative development of the transcription of all components involved, from the enzymes that drive denitrification to the anaerobic synthesis of the electron carrier ubiquinone and the O_2_-independent synthesis of the cofactor PQQ, which is essential for methanol dehydrogenase. The regulation of sulfur oxidation and anaerobic respiration are thus deeply intertwined, an aspect that, to our knowledge, has not yet been reported for chemoorganoheterotrophic sulfur oxidizers and has only become clear through the investigation of our *H. denitrificans* mutant strains.

In 1999, Sorokin provided the first direct evidence of the ability of obligately heterotrophic bacteria to oxidize thiosulfate under anaerobic conditions (Sorokin et al., 1999). The organisms investigated in that study cannot oxidize thiosulfate completely to sulfate but form tetrathionate as an end product, while the *H. denitrificans* X^T^ wildtype strain can pursue both pathways. The reference strain we investigated in the current study exclusively produces sulfate. The free energy released by reaction of thiosulfate with oxygen or nitrate is in a very similar range, with the process being sightly less favorable under anaerobic conditions [equations 1 and 2], especially when considering the more reduced state of the respiratory chain in the absence of oxygen (Sorokin et al., 1999).

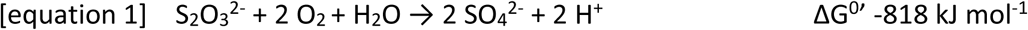

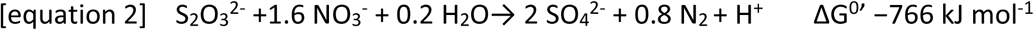

Heterotrophic thiosulfate-oxidizing nitrate reducers are vital in ecosystems where oxygen is limited, such as in hypoxic or anoxic zones. In marine sediments, wastewater treatment systems, and other anoxic environments, these bacteria contribute to the cycling of nitrogen and sulfur, affecting the overall health and functioning of these systems. Their ability to oxidize sulfur compounds and reduce nitrate makes them important contributors to the nitrogen and sulfur cycles. Among inorganic electron donors for nitrate reduction that may typically occur in subsurface environments, thiosulfate is a both readily available and non-toxic (Cardoso et al., 2006; Zhu and Getting, 2012; Kumar et al., 2018). In addition, several studies have investigated thiosulfate as an electron donor for nitrate removal in wastewater treatment but in depth studies of control mechanisms have not been performed (Cardoso et al., 2006). Knowledge concerning growth of *H. denitrificans* in the simultaneous presence of thiosulfate and nitrate is not available. It has been reported that the organism is able of aerobic denitrification in the presence of methylated amines (Meiberg et al., 1980), whereas on methanol nitrate reduction was not observed when cultures were incubated with oxygen and nitrate at the same time (Martineau et al., 2015).

## CONCLUSIONS

Our work much expands the role of the sulfane-sulfur responsive regulators sHdr and SoxR, provides evidence that their cooperative action, possibly even heterodimer formation, is indispensable for proper function *in vivo*, and underscores the notion that their activity also includes interaction with other transcriptional regulators. Most importantly, our work expands the role of the sHdrR/SoxR regulatory system far beyond sulfur oxidation and shows that a profound effect is exerted on anaerobic metabolism, in particular denitrification in *H. denitrificans*. The present study has thus set the stage for future research, which will further elucidate the intricate relationship between oxidative sulfur metabolism and denitrification. Enhanced understanding of these processes promises significant insights into the biology of these bacteria, particularly their role in environmental contexts, such as their contribution to greenhouse gas emissions (e.g., N₂O).

## Supporting information

Table with RNAseq data

Tree sHdrR and relatives Newick format

## ETHICS STATEMENT

This study adheres to the ethical standards set by *Environmental Microbiology*. All experimental protocols involving environmental samples, microbial cultures, and data collection were conducted following ethical guidelines and with proper approval from relevant institutional or governmental bodies. The research respects the principles of scientific integrity, transparency, and reproducibility. No conflicts of interest were declared by the authors. The study ensures that all findings are presented truthfully, with proper citations to prior works, and all sources of funding are disclosed.

## DATA AVAILABILITY STATEMENT

RNA-Seq raw data files and processed data files are available via the NCBI GEO repository (accession number GSE282994)

## AUTHOR CONTRIBUTIONS

**Jinjing Li:** investigation, formal analysis; writing – review and editing. **Nora Schmitte:** investigation. **Kaya Törkel:** investigation. **Christiane Dahl:** Conceptualization, writing – review and editing, supervision, funding acquisition

## ACKNOWLEDGEMENTS

We thank Stefania de Benedetti for support with RNA isolation and Angelina Hallik for help with EMSA experiments.

## CONFLICT OF INTEREST STATEMENT

The authors declare no conflict of interest.

## FUNDING

This research was in part funded by German Science foundation, grant numbers Da 351/13-1, Da 351/14-1 and Da 351/8-2. J.L. was financed by a Scholarship of the China Scholarship Council.

## Notes

https://www.ncbi.nlm.nih.gov/geo/query/acc.cgi?acc=GSE282994

https://www.ncbi.nlm.nih.gov/geo/query/acc.cgi?acc=GSE278992

### Competing Interest Statement

The authors have declared no competing interest.

