## Supplementary figures and tables for "In *Hyphomicrobium denitrificans* two related sulfane-sulfur responsive transcriptional repressors regulate thiosulfate oxidation and have a deep impact on nitrate respiration and anaerobic biosyntheses"

**Supplementary Fig. 1:** Growth of *H. denitrificans* reference and mutant strains on methanol

**Supplementary Fig. 2:** Growth and thiosulfate consumption of *H. denitrificans* Δ*tsdA* carrying a *shdrR* complementation *in cis*.

**Supplementary Fig. 3.** Volcano plots of differentially expressed genes for the *H. denitrificans* strains Δ*tsdA*, Δ*tsdA* Δ*shdrR* and Δ*tsdA* Δ*soxR*.

Transcript abundance changes of genes encoding transcriptional regulators and neighboring genes from *Hyphomicrobium denitrificans* Δ*tsdA*

**Supplementary Fig. 4.** Amino acid sequence alignment of selected LysR-type regulators.

**Supplementary Table 1:** Strains, primers and plasmids

**Supplementary Table 2:** Occurrence of sHdrR-related proteins with two conserved cysteines ($Cys^{50}$ and $Cys^{116}$ in *Hd*sHdrR)

**Supplementary Table 3:** mRNAseq analysis of *H. denitrificans* strains *ΔtsdA ΔsoxR* and *ΔtsdA ΔshdrR,* part 1.

**Supplementary Table 4:** mRNAseq analysis of *H. denitrificans* strains Δ*tsdA* Δ*soxR* and Δ*tsdA* Δ*shdrR,* part 2.

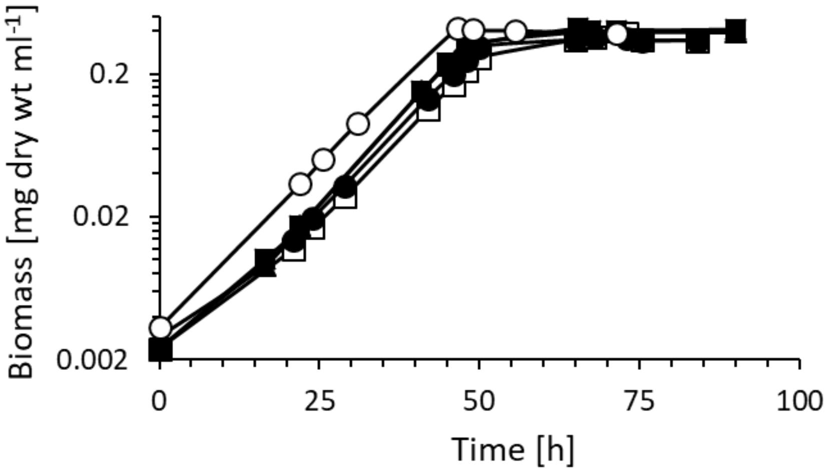


**Supplementary Fig. 1. Growth of *H. denitrificans* reference and mutant strains on methanol. a.** Growth curves are compared for the reference strain *H. denitrificans* Δ*tsdA* (filled circles), Δ*tsdA* Δ*shdrR* (open boxes), Δ*tsdA* sHdrR $C^{50}S$ (filled boxes), Δ*tsdA* sHdrR $C^{116}S$ (filled triangles), and Δ*tsdA* sHdrR $C^{50}S$ $C^{116}S$ (open circles). Error bars indicating SD for three replicates are too small to be visible for the determination of biomass.

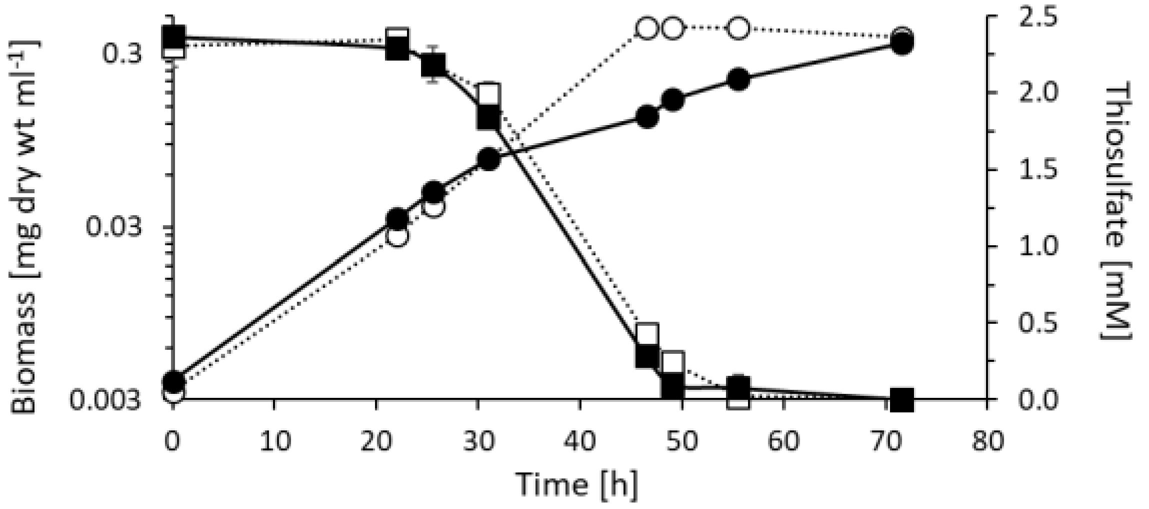


**Supplementary Fig. 2. Growth and thiosulfate consumption of *H. denitrificans* ΔtsdA carrying a *shdrR* complementation *in cis*.** Growth curves are shown for medium containing 2 mM thiosulfate. Pre-cultures were either thiosulfate-free (broken lines, open symbols) or were pre-induced and contained 2 mM thiosulfate (solid lines, filled symbols). Values for biomass and thiosulfate are given as circles and boxes, respectively. Error bars indicating SD for three replicates are too small to be visible for the determination of biomass.

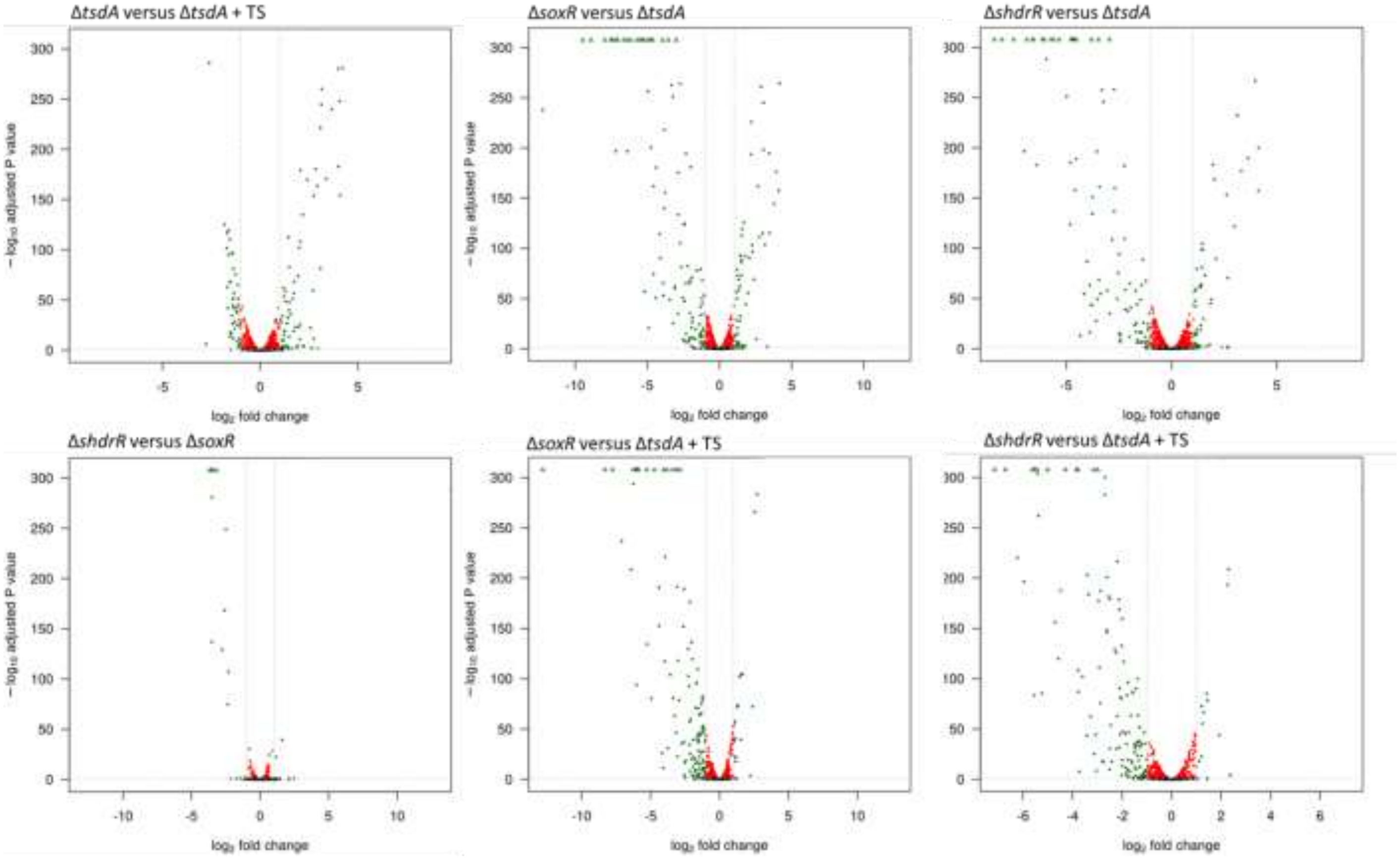


**Supplementary Fig. 3. Volcano plots of differentially expressed genes for the *H. denitrificans* strains Δ*tsdA*, Δ*tsdA* Δ*shdrR* and Δ*tsdA* Δ*soxR*.** The strains/conditions compared are given on top of each panel. TS, thiosulfate. • significant and has >1 log2-fold change, • significiant (FDR corrected p-value <= 0.1 • not significant.

```
CcmR            MGHHHHHHMQATLHQLKVFEATARHGSFTRAAEELYITQPTVSSQIKQLSKTVGLPLFEQ 60
YeiE            -----------TLRQLEVFAEVLKSGSTTQASQMLSLSQSAVSAALTDLEGQLGVQLFDR 49
Hden_0835       ----------MTLEQLRIFVAVAEREHVTQAAKELNLTQSATSAAVSALEARYATKLFDR 50
                           **.**.:*  . .    *:*:: * ::* :.*: :. *.   .  **::

CcmR            IGKRLYLTEAGQELLVTCQDIFQRLDNFAMKVADIKGTKQGRLRLAVI-TTAKYFIPRLL 119
YeiE            VGKRLVVNEHGRLLYPRTVALLEQAGEIERL----FRNDNGAIRVYASSTIGNYILPEII 105
Hden_0835       IGRRIVLTQAGKLFLVEAKSVLARAAAAEKVLADLAGLERGSLRIGASQTAGNYWLPEII 110
                :*:*: :.: *: :      :: :              ..* :*: .  * .:* :*.::

CcmR            GEFIQKYPGIEVSLKVTNHEQIRHRMQNNEDDLYIVSEPPEEIDLNYQPFLDNPLVVIAR 179
YeiE            ARYRRDFPDLPLEMSVGNSLDVVQAVCDFRVDIGLIEGPCHMAEIVAQPWLEDELVVFAS 165
Hden_0835       HRYQSLFPGISIALKIGNTETVAADVEDGVADLGFIEGEIDNPVLSVTPVADDDMVLVVA 170
                 .:   :*.: : :.: *   :   : :   *: ::.   .   :   *  :: :*:..

CcmR            RDHPLAGKSNIPITALNDEAFIMREKGSGTRLAVQNLFHR---HYVDVRVRLELGSNEAI 236
YeiE            PASPLLEGEV-TLERLAAMPWILREKGSGTREIVDYLLLS---HLPQFRLSMELGNSEAI 221
Hden_0835       PNNPLAKQPLRALSQIAQARWVVREAGSGTRAILEADVAKLGIDPKSLDIALELPSNEAV 230
                   **       :  :    :::** *****  ::  .     .  .. : :** ..**:

CcmR            KQAIAGGMGISVLSQHTLVSEGARSELTILDIDEFPIKRRWYVANLAGKQLSVITQTFLD 296
YeiE            KHAVRHGLGVSCLSRRVIAEQLETGSLVEVKVPLPPLVRTLYRIHHRQKHLSSALARFLR 281
Hden_0835       RGAVVAGSGITILSRLVVAAPLKAKTLVALDVPLPAR--KFFALRHKERYFTRAERTFID 288
                : *:  * *:: **: .:.       *. :.:         :  .   : ::     *:

CcmR            YLMAVTKNMPAPFAEQLTTQQTPVKLVL        324
YeiE            YCEL---------------------------        285
Hden_0835       VATGKQSSRAPG-------------------        300
```

**Supplementary Fig. 4. Amino acid sequence alignment of selected LysR-type regulators.** The protein deduced from Hden_0635 is aligned with CcmR/NdhR from *Synechocystis* PCC6803 (PDB 5Y2V) and YeiE from *Cronobacter sakazakii* (GenBank accession number ELY4740156). For *Synechocystis* CcmR/NdhR the residues interacting with 2-phosphoglycolate, which is an inducer (Jiang et al., 2018), are highlighted in green. For YeiE the residues interacting with sulfite (Hong et al., 2022) are marked yellow. An * (asterisk) indicates positions with identical residues. Cysteines are highlighted in yellow. Colons (:) and single dots (.) indicate conserved and semi-conserved amino acids, respectively.

### Supplementary Table 1. Strains, primers and plasmids

| Strains primers or plasmids | Relevant genotype, description or sequence | Reference or source |
|---|---|---|
| **Strains** | | |
| *Escherichia coli* 10-beta | Δ*(ara-leu) 7697 araD139 fhuA* Δ*lacX74 galK16 galE15 e14-* Φ*80*d*lacZ*Δ*M15 recA1 relA1 endA1 nupG rpsL* ($Str^R$) *rph spoT1* Δ*(mrr-hsdRMS-mcrBC)* | New England Biolabs |
| *E. coli* DH5α | $F^-$ Φ80*lac*ZΔM15 Δ(*lac*ZYA-*arg*F)U169 *rec*A1 *end*A1 *hsd*R17($r_K^-$, $m_K^+$) *pho*A *sup*E44 $\lambda^-$*thi*-1 *gyr*A96 *rel*A1 | New England Biolabs |
| *E. coli* BL21(DE3) | $F^-$*omp*T *hsd*$S_B$ ($r_B^-$, $m_B^-$) *gal dcm* (DE3) | Novagen |
| *Hyphomicrobium denitrificans* Δ*tsdA* | $Sm^R$, in-frame deletion of *tsdA* in *H. denitrificans* Sm200 | (Koch and Dahl, 2018) |
| *H. denitrificans* Δ*tsdA* Δ*shdrR* | $Sm^R$, in-frame deletion of *shdrR* (Hden_0682) in *H. denitrificans* Δ*tsdA* | (Li et al., 2023b) |
| *H. denitrificans* Δ*tsdA* Δ*soxR* | $Sm^R$, deletion of *soxR* (Hden_0700) in *H. denitrificans* Δ*tsdA* | (Li et al., 2023a) |
| *H. denitrificans* Δ*tsdA shdrRcomp* | $Sm^R$, *cis* complementation of *H. denitrificans* Δ*tsdA* Δ*shdrR* with *shdrR* | This work |
| *H. denitrificans* Δ*tsdA shdrR-Cys*$^{50}$*Ser* | Exchange of sHdrR-Cys$^{50}$ to Ser in *H. denitrificans* Δ*tsdA* | This work |
| *H. denitrificans* Δ*tsdA shdrR-Cys*$^{116}$*Ser* | Exchange of sHdrR-Cys$^{116}$ to Ser in *H. denitrificans* Δ*tsdA* | This work |
| *H. denitrificans* Δ*tsdA shdrR- Cys*$^{50}$*Ser-Cys*$^{116}$*Ser* | Exchange of sHdrR- Cys$^{50}$ and Cys$^{116}$ to Ser in *H. denitrificans* Δ*tsdA* | This work |
| **Primers** | | |
| Fr-pET22b-sHdrR-trun-NdeI | GGCA**CATATG**ACCGACGCGTCGATCGAACAG (NdeI) | This work |
| Rev-pET22b-0682-NotI | TTTT**GCGGCCGC**ATTCGAGCGTTTTCCCGCAC (NotI) | (Li et al., 2023b) |
| sHdrR _C50S_Up_Rev | GGTTCTTTCTCCCTCGAGCAGGAGGGACAAAATCGCGAGA | This work |
| sHdrR _C50S_Down_Fw | TCTCGCGATTTTGTCCCTCCTGCTCGAGGGAGAAAGAACC | This work |
| sHdrR _C116S _Up_rev | GTTTTCCCGCACTCGTTGCACTATAATACTTATGCAGCGT | This work |
| sHdrR_C116S_Down_Fw | ACGCTGCATAAGTATTATAGTGCAACGAGTGCGGGAAAAC | This work |
| Fwd_deltaHden0682_BamHI | GCAT**GGATCC**GCGAAAATGTGCACCGGAG (BamHI) | (Li et al., 2023b) |
| Rev_deltaHden0682_XbaI | AAGC**TCTAGA**TATGCGGCAGCCGTTGACGC (XbaI) | (Li et al., 2023b) |
| EMSA-Fr | TTCCCGCCCCGTCTTGGTTT | (Li et al., 2023b) |
| EMSA_Fr2_Fr | TCAGCGCTCGCCTGGAAGTC | (Li et al., 2024) |
| EMSA_Fr3_Rev | TCTAAGCATCAACATATTCATATCTTTATATATTTTCG | (Li et al., 2024) |
| EMSA-Rev | AGGAGTTGCATCCAAAAAAGCGTG | (Li et al., 2023b) |
| EMSA-Hden_0703/04-fw | GGGTCACCAAATTCTGCAGGTCTC | (Li et al., 2024) |
| EMSA-Hden_0703/04-rev | ATCACGCCATCTCTCCCGGAA | (Li et al., 2024) |
| EMSA-Hden_0699/0698-fw | AATTCCACGGCTCCGCC | (Li et al., 2024) |
| EMSA-Hden_0699/0698-rev | TCGACAGCTTGCGGAAATCC | (Li et al., 2024) |
| EMSA-sHdrR-LipS1_F | TAGAGCGAGTCTTCAGC | (Li et al., 2024) |
| EMSA-sHdrR-LipS1_R | CGGCCCTCTGAGAAAAG | (Li et al., 2024) |
| EMSA-LipX-DsrE_F | GACTTCGCCGATCAATCGATC | (Li et al., 2024) |
| EMSA-LipX-DsrE_R | TGCCACCTCCCCGATATG | (Li et al., 2024) |
| EMSA-Hden_0703/04-fw | GGGTCACCAAATTCTGCAGGTCTC | (Li et al., 2024) |
| rpoB-denitf | AGGACGTGTTCACCTCGATT | (Martineau et al., 2015) |
| rpoB-denitr | CGGCTTCGTCAAGGTTCTTC | (Martineau et al., 2015) |
| SoxT1A 0681_qPCR-Fr | CCCGAGTGATACGATTCGCA | (Li et al., 2023a) |
| SoxT1A 0681_qPCR-Rev | CTAAAATGCCGCCGGTGATG | (Li et al., 2023a) |
| LplA_qPCR-Fr | GGCCATGATCGATTTGCACC | (Li et al., 2024) |
| LplA_qPCR-Rev | CGAGATAAATTGCACCGCCG | (Li et al., 2024) |
| sHdrA_qPCR-Fr | CCGATCACCATTCCGTTCGA | (Li et al., 2023a) |
| sHdrA_qPCR-Rev | CAATTGTTTCCGGGCCGATC | (Li et al., 2023a) |
| sHdrB2_qPCR-Fr | GACGTGGCCTACTATTCCGG | (Li et al., 2024) |
| sHdrB2_qPCR-Rev | CCGCGACGACAGATAGGTTT | (Li et al., 2024) |
| LbpA2_qPCR-Fr | GGTTCCAAGAGCAGCCTGAT | (Li et al., 2024) |
| LbpA2_qPCR-Rev | TCGTTGATCTCCAGAACCGC | (Li et al., 2024) |
| SoxXA_qPCR-Fr | CGGCGCTCATTACCTATCTC | (Li et al., 2024) |

| | | |
|---|---|---|
| SoxXA_qPCR-Rev | TCGGGGTGTCTTTTTCAGTC | (Li et al., 2024) |
| TusA_qPCR-Fr | TCTGACAGTTGATGCCAAGG | (Li et al., 2024) |
| TusA_qPCR-Rev | CGTTTCCTCATGTTCAAGCA | (Li et al., 2024) |
| CytP450_qPCR-Fr | CAATACGGTTCTCGGACGTT | (Li et al., 2024) |
| CytP450_qPCR-Rev | CATTCGTTTCCTGACGAGGT | (Li et al., 2024) |
| SoxT1B (0699)_qPCR-Fr | GCCGCCGTCTCAGTAAATAA | (Li et al., 2024) |
| SoxT1B (0699)_qPCR-Rev | AGCAGAAGACGGCAGATGAT | (Li et al., 2024) |
| SoxR_qPCR-Fr | TGAAGCGGACGAGGAAGTAT | (Li et al., 2024) |
| SoxR_qPCR-Rev | GAGACTGTGGGCTGGTTGAT | (Li et al., 2024) |
| sHdrR_qPCR-Fr | TTAGGAAGTCCGCATCGTCT | (Li et al., 2024) |
| sHdrR_qPCR-Rev | GCACTCGTTGCGCAATAATA | (Li et al., 2024) |
| SoxY_qPCR-Fr | GTTCAGCTTGCGGACTTTTC | (Li et al., 2024) |
| SoxY_qPCR-Rev | GCCAATCGTCACCTTCACTT | (Li et al., 2024) |
| **Plasmids** | | |
| pHP45Ω-Tc | $Ap^r$, $Tc^r$ | (Fellay et al., 1987) |
| pk18*mobsacB* | $Km^r$, $Mob^+$, *sacB*, *oriV*, *oriT*, *lacZα* | (Schäfer et al., 1994) |
| pET-22b (+) | ApR, T7 promoter, lac operator, C-terminal His tag, pelB leader | Novagen |
| pET-22bHdsHdrR-trunc | $Ap^R$, NdeI-NotI fragment of PCR amplified truncated *shdrR* in NdeI-NotI of p ET-22b (+) | This work |
| pk18*mobsacB-shdrR* | $Km^r$, 2379 bp PCR fragment for chromosomal complementation of *shdrR* cloned int pk18*mobsacB* using XbaI and BamHI sites | This work |
| pk18*mobsacB-shdrR*-Tc | $Km^r$, $Tc^r$, pHP45ΩTc tetracycline cassette inserted into pk18*mobsacB-shdrR* using SmaI | This work |
| pk18*mobsacB-shdrR-C50S* | $Km^r$, SOE PCR fragment implementing chromosomal integration of *shdrR* encoding a $Cys^{50}$Ser exchange cloned into pk18*mobsacB* using XbaI and BamHI restriction sites | This work |
| pk18*mobsacB-shdrR-C50S*-Tc | $Km^r$, $Tc^r$, pHP45ΩTc tetracycline cassette inserted into pk18*mobsacB-shdrR-C50S* using SmaI | This work |
| pk18*mobsacB-shdrR-C116S* | $Km^r$, SOE PCR fragment implementing chromosomal integration of *shdrR* encoding a $Cys^{116}$Ser exchange cloned into pk18*mobsacB* using XbaI and BamHI restriction sites | This work |
| pk18*mobsacB-shdrR-C116S*-Tc | $Km^r$, $Tc^r$, pHP45ΩTc tetracycline cassette inserted into pk18*mobsacB-shdrR-C116S* using SmaI | This work |
| pk18*mobsacB-shdrR-C50S-C116S* | $Km^r$, SOE PCR fragment implementing chromosomal integration of *shdrR* encoding $Cys^{50}$Ser and $Cys^{116}$Ser exchange cloned into pk18*mobsacB* using XbaI and BamHI restriction sites | This work |
| pk18*mobsacB-shdrR-C50S-C116S*-Tc | $Km^r$, $Tc^r$, pHP45ΩTc tetracycline cassette inserted into pk18*mobsacB-shdrR-C50S-C116S* using SmaI | This work |

**Supplementary Table 2. Occurrence of sHdrR-related proteins with two conserved cysteines (Cys[50] and Cys[116] in *Hds*HdrR).** Accession number and/or locus tags are provided. Linked genes were manually analyzed. Furthermore, genomes were checked via HMSS2 (Tanabe and Dahl, 2023) for the presence of genes encoding Sox-dependent thiosulfate oxidation in the periplasm (set positive when *soxYZAXB* were detected, thus covering complete and truncated Sox systems (Li et al., 2023a)) and sHdr-driven sulfane sulfur oxidation in the cytoplasm (set positive when at least 70 % of the genes *shdrC1B1AHC2B2* or *shdrC1B1AHB3etfAB* were present in a syntenic block, respectively (Kümpel et al., 2024)).

| Organism | Accession, locus tag | Linked genes | sHdr system | Sox system | References |
|---|---|---|---|---|---|
| **Pseudomonadota** | | | | | |
| **Alphaproteobacteria** | | | | | |
| **Hyphomicrobiales** | | | | | |
| ***Hyphomicrobiaceae*** | | | | | |
| *Hyphomicrobium denitrificans* X$^T$ (ATCC 51888$^T$) | sHdrR: Hden_0682<br>SoxR: Hden_0700 | *shdr-lbpA*<br>*sox* | Yes | Yes | |
| *Hyphomicrobium denitrificans* 1NES1 | HYPDE_25308 | RND transporter | No | No | (Venkatramanan et al., 2013) |
| *Hyphomicrobium* sp. GJ21 | sHdrR: HYPGJ_30422<br>SoxR: HYPGJ_30404 | *shdr-lbpA*<br>*sox* | Yes | Yes | (Tatusova et al., 2014) |
| *Hyphomicrobium* sp. SCN 65-11 | ABS54_17655 | Short fragment | No | No | (Kantor et al., 2015) |
| *Hyphomicrobium* sp. CS1BSMeth3 | WP_083528837: CS1BSM3_04686<br>WP_210188842: CS1BSM3_RS16485 | TauE, *ccm* genes<br>RND transporter | Yes | Yes | Adelskov and Patel, unpublished |
| *Hyphomicrobium* sp. FW.3.32 | CTY20_06775 | *soxBZYAX* | No | Yes | (Zhang et al., 2017) |
| *Filomicrobium insigne* CGMCC 1.6497$^T$ | SAMN04488061_2704<br>SoxR: SAMN04488061_1979 | Only *soxYZ*<br>*soxCBZY* | No | Yes | (Wu et al., 2009) |
| *Rhodomicrobium vaniielii* ATCC 17100$^T$ | MBJ7534237 JDN40_08990<br>MBJ7535956 JDN40_17755 | *tauE*<br>RND transporter | No | SoxXA present | Conners et al, unpublished |
| ***Devosiaceae*** | | | | | |
| *Devosia nanyangense* NC_groundwater_1586_Pr3_B-0.1um_66_15 | HY834_20740 | *shdr* gene cluster, *tusA* | Yes | No | (He et al., 2021) |
| **Acetobacterales** | | | | | |
| ***Acetobacteriaceae*** | | | | | |
| *Rhodopila globiformis* DSM 161$^T$ | CCS01_RS26760<br>CCS01_RS13140 | RND transporter, Rhd<br>*shdr* genes | Yes | No | (Imhoff et al., 2018) |
| **Rhizobiales** | | | | | |
| ***Rhizobiaceae*** | | | | | |
| *Agrobacterium fabrum (tumefaciens*) C58$^T$ | BIGR_AGRFC, Atu3466 | Rhd-PDO fusion-*bigR-pmpBA* | No | No | (Goodner et al., 2001; Guimarães et al., 2011) |
| *Pseudaminobacter salicylatoxidans* KCT001 | WP_019171658 | *sox* | No | Yes | (Mandal et al., 2007) |

| | | | | | |
|---|---|---|---|---|---|
| ***Xanthobacteraceae*** | | | | | |
| *Bradyrhizobium diazoefficiens* USDA10$^T$ | BAC48771 | *sox* | No | Yes | (Kaneko et al., 2002) |
| *Rhodopseudomonas palustris* TIE-1 | Rpal_4967 | Between *sox* genes and genes for RND transporter | No | Yes | (Larimer et al., 2004) |
| *Rhodoplanes elegans* DSM 11907$^T$ | RAI38494, CH338_12525 | RND transporter, sulfurtransferase | No | Yes | (LaSarre et al., 2018) |
| **Rhodospirillales** | | | | | |
| ***Rhodospirillaceae*** | | | | | |
| *Rhodospirillum rubrum* ATCC 11170$^T$ | WP_011389407, ABC22517, Rru_A1717 | *pmpB-like* | No | No | (Munk et al., 2011) |
| **Rhodobacterales** | | | | | |
| ***Paracoccaceae*** | | | | | |
| *Paracoccus denitrificans* GB17 | CAB94376 | *sox* genes | No | Yes | (Wodara et al., 1997; Rother et al., 2005) |
| ***Rhodobacteraceae*** | | | | | |
| *Roseobacter litoralis* Och 149$^T$ | AEI95148 | *sox* genes | No | Yes | (Kalhoefer et al., 2011) |
| *Rhodobacter capsulatus* SB 1003 | ADE85198 | RND transporter | No | No | (Shimizu et al., 2017; Capdevila et al., 2021) |
| *Rhodovulum sulfidophilum* DSM 1374$^T$ | AAO11780 | *sox* genes | No | Yes | (Appia-Ayme et al., 2001) |
| **Sphingomonadales** | | | | | |
| ***Sphingomonadaceae*** | | | | | |
| *Tsuneonella (Altererythrobacter) mangrovi* CD9-11$^T$ | WP_240504499: CJO11_RS12710 | close to *shdr* genes and genes for RND transporter | Yes | No | (Tatusova et al., 2014) |
| *Erythrobacter* sp. NAP1 | EAQ29854, NAP1_03740 | RND efflux system | No | | (Koblizek et al., 2011) |
| **Gammaproteobacteria** | | | | | |
| **Chromatiales** | | | | | |
| ***Chromatiaceae*** | | | | | |
| *Allochromatium vinosum* DSM 180$^T$ | Alvin_3027 | Rhd | No | Yes | (Weissgerber et al., 2011) |
| **Nitrococcales** | | | | | |
| ***Ectothiorhodospiraceae*** | | | | | |
| *Halorhodospira* halophila DSM244$^T$ | Hhal_1425 | RND transporter | No | Yes | (Challacombe et al., 2013) |
| **Enterobacterales** | | | | | |
| ***Enterobacteriaceae*** | | | | | |
| *Escherichia coli* O1 strain PSU-0611 | EEZ6061186, DCO30_005030 | Short fragment | No | No | (Lacher et al., 2020) |
| *Escherichia coli* K12 substr. MG1655 | b2667; YgaV PDB: 3CUO | YgaP: membrane-associated protein with rhodanese activity | No | No | (Paul and Larson, 2006; Riley et al., 2006; Gueuné et al., 2008) |

| | | | | | |
|---|---|---|---|---|---|
| ***Vibrionaceae*** | | | | | |
| *Vibrio cholera* O1 biovar El Tor N16961 | HlyU, VC_0678, HLYU_VIBCH, PDB: 4K2E<br>VC_A0642, AAF96543 | Transcriptional activator of hemolysin, | No | No | (Williams et al., 1993; Heidelberg et al., 2000; Mukherjee et al., 2014) |
| **Xanthomonadales** | | | | | |
| ***Xanthomonadaceae*** | | | | | |
| *Xylella fastidiosa* 9a5c | WP_010893290, XF_0767, PDB: 3PQJ | Blh: DUF442-PDO fusion (XF_0768) *pmpBA* (XF_0765, 0766) | No | No | (Simpson et al., 2000; Barbosa and Benedetti, 2007; Guimarães et al., 2011) |
| **Burkholderiales** | | | | | |
| ***Chromobacteriaceae*** | | | | | |
| *Chromobacterium violaceum* ATCC 12472$^T$ | CV_0084 | *pmpAB,* Cyt c4 | No | No | (Brazilian National Genome Project, 2003) |
| ***Burkholderiaceae*** | | | | | |
| *Comamonas aquatica* CJG | WP_045267543 | *pmpAB,* DUF599 family | No | No | (Dai et al., 2016) |
| ***Thiobacillaceae*** | | | | | |
| *Thiobacillus denitrificans* ATCC 25No250 | AAZ98348, Tbd_2395 | alone | No | Yes | (Beller et al., 2006) |
| **Acidithioacillales** | | | | | |
| ***Acidithiobacillaceae*** | | | | | |
| *Acidithiobacillus thiooxidans* ATCC 19377$^T$ | WP_024893036.1, GCD22_RS14465 | RND transporter | Yes | Yes | (Valdes et al., 2011) |
| **Bacteroidota** | | | | | |
| **Bacteroidia** | | | | | |
| **Chitinophagales** | | | | | |
| Sphingobacteriales bacterium PMG_127 | RYD90138 | PmpA or B (short contig) | - | - | (Crombie et al., 2018) |
| **Bacillota** | | | | | |
| **Bacilli** | | | | | |
| **Lactobacillales** | | | | | |
| ***Streptpcoccaceae*** | | | | | |
| *Streptococcus pneumonia* SMRU2535 | CJK49847, ERS022045_00348 | *pmpBA*, uncharacterized, Rhd, sulfate permease, sulfide dehydrogenase | No | No | (Chewapreecha et al., 2014) |
| **Staphylococcales** | | | | | |
| ***Staphylococcaceae*** | | | | | |
| *Staphylococcus aureus* VB1919 | RTY94661 | Short contig | No | No | Balaji and Yamuna unpublished |

| | | | | | |
|---|---|---|---|---|---|
| **Clostridia** | | | | | |
| **Clostridiales** | | | | | |
| ***Clostridiaceae*** | | | | | |
| *Hathewaya proteolytica* DSM 3090$^T$ | SHJ53940, SAMN02745248_00335 | Upstream two genes for FeS containing proteins, then two for sulfur carrier protein ThiS | No | No | Joint Genome Institute |
| **Cyanobacteriota** | | | | | |
| **Cyanobacteriia** | | | | | |
| **Cyanobacteriales** | | | | | |
| ***Geitlerinemaceae*** | | | | | |
| *Sodalimena* (former *Phormidium*) *willei* BDU 130791 | OAB56254, AY600_15135 | All hypothetical | No | No | Peter. et al unpublished |

Ccm, cytochrome c maturation (Thöny-Meyer, 2002); PDO, persulfide dioxygenase; PmpAB, members of the YeeE/YedE family of transporters that have been predicted to transport sulfur-containing ions (Gristwood et al., 2011); Rhd, rhodanese; RND, Resistance-nodulation-division family transporters, a category of bacterial efflux pumps, especially identified in Gram-negative bacteria (Nikaido, 2011); TauE, sulfite exporter (Weinitschke et al., 2007)

**Supplementary Table 3. mRNAseq analysis of *H. denitrificans* strains *ΔtsdA ΔsoxR* and *ΔtsdA ΔshdrR*, part 1.** Genes with higher mRNA abundance in the regulator-deficient mutants than in the reference strain in the absence of thiosulfate.

| Locus tag | Annotation[a] | ***ΔtsdA ΔsoxR* vs *ΔtsdA*** Fold change** | ***ΔtsdA ΔshdrR* vs *ΔtsdA*** Fold change** |
|---|---|---|---|
| **Sulfur metabolism** | | | |
| Hden_0678 | hypothetical protein | 5.82 | 3.51 |
| Hden_0679 | DsbA family protein | 11.06 | 2.17 |
| Hden_0680 | Rhd, sulfur transferase domain-containing protein | 15.61 | 2.60 |
| Hden_0681 | SoxT1A, YeeE/YedE family protein | 24.43 | 4.32 |
| Hden_0683 | LipS1, radical SAM protein | 17.88 | 25.64 |
| Hden_0684 | LipT, NAD(P)/FAD-dependent oxidoreductase | 17.27 | 17.42 |
| Hden_0685 | LipS2, radical SAM protein | 14.96 | 15.70 |
| Hden_0686 | Lpl(AB), lipoate--protein ligase family protein | 16.04 | 18.56 |
| Hden_0687 | LipX, GMP synthase - glutamine amidotransferase domain-like protein | 15.25 | 17.49 |
| Hden_0688 | DsrE3C, DsrE/DsrF/DrsH-like family protein | 10.84 | 15.72 |
| Hden_0689 | sHdrC1 | 8.19 | 11.79 |
| Hden_0690 | sHdrB1 | 7.40 | 10.69 |
| Hden_0691 | sHdrA, FAD-dependent oxidoreductase | 8.21 | 11.65 |
| Hden_0692 | sHdrH | 7.93 | 12.36 |
| Hden_0693 | sHdrC2 | 6.72 | 9.96 |
| Hden_0694 | sHdrB2 | 6.37 | 8.71 |
| Hden_0695 | sHdrI | 5.24 | 7.90 |
| Hden_0696 | LbpA2 | 4.17 | 6.12 |
| Hden_0697 | Cytochrome P450 | 4.77 | ns |
| Hden_0698 | TusA family sulfurtransferase | 4.88 | ns |
| Hden_0699 | SoxT1B, YeeE/YedE family protein | 2.69 | ns |
| Hden_0701 | SoxS, thioredoxin family protein | 13.69 | ns |
| Hden_0702 | Sulfur oxidation c-type cytochrome SoxX | 12.43 | ns |
| Hden_0703 | Sulfur oxidation c-type cytochrome SoxA | 17.19 | ns |
| Hden_0704 | Thiosulfate oxidation carrier protein SoxY | 17.44 | ns |
| Hden_0705 | Thiosulfate oxidation carrier complex protein SoxZ | 16.72 | ns |
| Hden_0706 | Thiosulfohydrolase SoxB | 14.50 | ns |
| Hden_0834 | YeiH family protein, sulfite export | ns | 3.70 |
| **Carbon metabolism** | | | |
| Hden_0802 | DUF3734 domain-containing protein | 2.42 | ns |
| Hden_2747 | Acyl CoA:acetate/3-ketoacid CoA transferase | 2.33 | 2.12 |
| **Heme degradation and iron acquisition** | | | |
| Hden_0540 | TonB-dependent heme receptor | 4.61 | ns |
| Hden_0541 | Heme degrading monooxygenase HmoA | 3.45 | 3.02 |
| Hden_0542 | TonB family protein | 3.20 | 2.82 |
| Hden_0874 | Hemin uptake protein HemP | 3.53 | 3.58 |
| Hden_0875 | Heme degrading monooxygenase HmoA | 2.86 | 2.62 |
| Hden_0876 | Heme utilization cystosolic carrier protein ChuX/HutX | 2.88 | 2.69 |
| Hden_0877 | Heme transport system substrate-binding protein ChuT | 2.97 | 2.77 |
| Hden_0878 | Heme transport system permease protein ChuU | 2.82 | 2.74 |
| Hden_0879 | Heme transport system ATP-binding protein, HmuV | 2.64 | 2.68 |

| | | | |
|---|---|---|---|
| Hden_1331 | TonB-dependent siderophore receptor | 2.54 | 2.24 |
| Hden_1332 | PepSY domain-containing protein | 2.23 | ns |
| Hden_1333 | hypothetical protein | 2.25 | ns |
| Hden_3200 | hypothetical protein | 10.51 | 8.75 |
| Hden_3201 | hypothetical protein | 9.99 | 7.27 |
| Hden_3202 | Hemin uptake protein HemP | 8.85 | 6.35 |
| **Transport** | | | |
| Hden_0532 | ABC transporter substrate-binding protein, branched chain amino acid transport | 2.73 | ns |
| Hden_2931 | Potassium-transporting ATPase subunit KdpA | 2.92 | 2.08 |
| Hden_3198 | YceI family protein, periplasmic, polyisoprenoid-binding | 2.23 | 2.04 |
| Hden_3199 | Cytochrome *b*, YceJ | 2.64 | 2.31 |
| **Regulation** | | | |
| Hden_0594 | helix-turn-helix domain-containing protein | 2.60 | 2.39 |
| Hden_0722 | response regulator transcription factor, LuxR family | 2.58 | |
| Hden_2164 | AraC family transcriptional regulator | 3.05 | ns |
| **Respiration and electron transport** | | | |
| Hden_2084 | pseudoazurin | 4.56 | 4.11 |
| Hden_3539 | cupredoxin domain-containing protein | 2.40 | 2.12 |
| Hden_2748 | c-type cytochrome | 3.50 | 3.17 |
| Hden_2908 | cytochrome c oxidase subunit II | 2.32 | 2.33 |
| **Other** | | | |
| Hden_0136 | hypothetical protein | 1.00 | 2.27 |
| Hden_0441 | glycosyltransferase | 1.00 | 6.50 |
| Hden_0457 | hypothetical protein | 2.29 | 2.30 |
| Hden_0523 | zf-HC2 domain-containing protein | ns | 2.25 |
| Hden_0525 | catalase family peroxidase | ns | 3.92 |
| Hden_0738 | hypothetical protein | 2.18 | 2.23 |
| Hden_0914 | hypothetical protein | ns | 5.23 |
| Hden_0990 | hypothetical protein | ns | 2.01 |
| Hden_1114 | hypothetical protein | 2.51 | 2.34 |
| Hden_1235 | phage GP46 family protein | ns | 2.06 |
| Hden_1416 | hypothetical protein | 2.01 | ns |
| Hden_1432 | DUF3307 domain-containing protein | ns | 2.26 |
| Hden_1518 | hypothetical protein | ns | 2.28 |
| Hden_2458 | hypothetical protein | 9.73 | 6.35 |
| Hden_2517 | hypothetical protein | 2.49 | ns |
| Hden_2518 | catalase | 2.85 | 2.73 |
| Hden_2542 | class I SAM-dependent methyltransferase | ns | 2.15 |
| Hden_2599 | hypothetical protein | 2.31 | 2.54 |
| Hden_2684 | hypothetical protein | ns | 2.30 |
| Hden_2944 | DUF3302 domain-containing protein | 2.01 | ns |
| Hden_2965 | hypothetical protein | ns | 2.41 |
| Hden_2982 | hypothetical protein | ns | 2.35 |
| Hden_3015 | DNA cytosine methyltransferase | ns | 2.22 |
| Hden_3020 | hypothetical protein | ns | 2.70 |
| Hden_3022 | hypothetical protein | 3.27 | ns |
| Hden_3142 | FHA domain-containing protein | 2.14 | 2.10 |
| Hden_3444 | hypothetical protein | ns | 2.19 |
| Hden_3518 | hypothetical protein | 2.54 | 2.18 |
| Hden_R0029 | | 2.00 | 2.21 |

| | | |
|---|---|---|
| Hden_R0052 | ns | 2.47 |

ns, not significant

[a] Gene names obtained using sequence similarities in Uniprot or NCBI databases

** Significance threshold set at >2-fold change and $p<0.001$;

**Supplementary Table 4. mRNAseq analysis of *H. denitrificans* strains Δ*tsdA* Δ*soxR* and Δ*tsdA* Δ*shdrR*, part 2.** Genes with lower mRNA abundance in the regulator-deficient mutants than in the reference strain in the absence of thiosulfate. Potential regulatory proteins associated with genes for respiratory proteins are printed in bold and not arranged under the headline "regulation".

| Locus tag | Annotation[a] | Δ*tsdA* Δ*soxR* vs. Δ*tsdA* Fold change** | Δ*tsdA* Δ*shdrR* vs. Δ*tsdA* Fold change** |
|---|---|---|---|
| **Biosynthesis of metabolites and cofactors** | | | |
| **PQQ** | | | |
| Hden_0547 | FmdE family protein, Flag1 repressor motif | 0.175 | 0.184 |
| Hden_0550 | urate hydroxylase PuuD | ns | 0.427 |
| Hden_0551 | pyrroloquinoline quinone biosynthesis protein PqqE | 0.424 | 0.362 |
| Hden_0552 | pyrroloquinoline quinone biosynthesis peptide chaperone PqqD | 0.190 | 0.185 |
| Hden_0553 | pyrroloquinoline quinone precursor peptide PqqA | 0.232 | 0.321 |
| **Fatty acids** | | | |
| Hden_0554 | beta-ketoacyl-ACP synthase FabF | 0.067 | 0.056 |
| Hden_0555 | beta-ketoacyl-ACP synthase FabF | 0.057 | 0.036 |
| Hden_0556 | zinc-binding dehydrogenase, putative enoyl-ACP reductase FabI function | 0.012 | 0.008 |
| Hden_0557 | beta-ketoacyl-ACP synthase FabF | 0.007 | 0.012 |
| Hden_0558 | beta-ketoacyl-ACP synthase FabF | 0.010 | 0.016 |
| Hden_0559 | beta-hydroxyacyl-ACP dehydratase FabZ | 0.004 | 0.006 |
| Hden_0560 | acyl carrier protein | 0.001 | 0.003 |
| Hden_0561 | 3-oxoacyl-ACP reductase, FabG | 0.002 | 0.004 |
| Hden_0562 | HAD-IIIC family phosphatase, putative involvement in methoxymalonyl-ACP biosynthesis, FkbH-like protein | 0.025 | 0.038 |
| Hden_0563 | acyl carrier protein | 0.199 | 0.218 |
| **Ubiquinone** | | | |
| Hden_0564 | UbiX family flavin prenyltransferase | 0.219 | 0.049 |
| Hden_0565 | UbiD family decarboxylase | 0.033 | 0.068 |
| Hden_0566 | UbiT ubiquinone biosynthesis accessory factor UbiT, SCP2 sterol-binding domain-containing protein | 0.046 | 0.071 |
| Hden_0567 | $O_2$-independent ubiquinone biosynthesis protein UbiU | 0.028 | 0.073 |
| Hden_0568 | $O_2$-independent ubiquinone biosynthesis protein UbiV | 0.107 | 0.084 |
| Hden_0569 | Cytochrome P450 | 0.185 | 0.168 |
| **Hden_0570** | **Crp/Fnr family transcriptional regulator** | 0.138 | 0.093 |
| Hden_0571 | DUF2478 domain-containing protein | 0.411 | 0.421 |
| **Respiration and electron transport** | | | |
| Hden_0508 | Cupin domain-containing protein | 0.173 | 0.238 |
| Hden_0509 | SPW repeat protein | 0.238 | 0.321 |
| Hden_0510 | Ferredoxin-NADP reductase | 0.200 | 0.211 |
| Hden_0572 | Hypothetical protein | 0.042 | 0.097 |
| Hden_0573 | 4Fe-4S binding protein | 0.047 | 0.036 |
| Hden_0574 | Periplasmic cupredoxin domain-containing protein | 0.032 | 0.032 |
| Hden_0579 | NorE | ns | 0.423 |
| Hden_0581 | Nitric oxide reductase subunit C, NorC | 0.066 | 0.068 |
| Hden_0582 | Nitric oxide reductase subunit B, NorB | 0.215 | 0.177 |
| Hden_0583 | Nitric oxide reductase NorQ protein | 0.453 | 0.396 |

| | | | |
|---|---|---|---|
| Hden_0584 | Nitric oxide reductase NorD protein, VWA domain-containing protein | 0.457 | 0.486 |
| Hden_0585 | Cytochrome c, hypothetical protein | 0.388 | 0.461 |
| Hden_0587 | DUF2946 domain-containing protein | 0.336 | 0.376 |
| Hden_0589 | Hypothetical protein | 0.137 | 0.131 |
| Hden_0590 | NnrS family protein, involved in response/tolerance to NO | 0.309 | 0.251 |
| Hden_0591 | Copper-containing nitrite reductase apoprotein NirK | 0.189 | 0.213 |
| Hden_0592 | Host attachment family protein | 0.042 | 0.042 |
| **Hden_0595** | **Helix-turn-helix domain-containing protein** | 0.037 | 0.043 |
| **Hden_0596** | **PAS domain-containing protein** | 0.259 | 0.309 |
| **Hden_0597** | **Signal transduction histidine kinase, nitrite/nitrate specific NarQ** | 0.073 | 0.074 |
| **Hden_0598** | **Two-component system response regulator NarL** | 0.059 | 0.062 |
| Hden_0673 | NitT/TauT family transport system substrate-binding protein, nitrate/sulfonate transport | 0.138 | 0.154 |
| Hden_0674 | NitT/TauT family transport system permease protein | 0.243 | 0.217 |
| Hden_0675 | NitT/TauT family transport system ATP-binding protein | 0.271 | 0.297 |
| Hden_0676 | NnrS family protein, involved in response/tolerance to NO | 0.317 | 0.349 |
| Hden_0677 | NnrS family protein, involved in response/tolerance to NO | 0.491 | 0.412 |
| Hden_0922 | VOC family protein | 0.201 | 0.326 |
| Hden_0924 | Porin | 0.086 | 0.090 |
| Hden_0925 | NarK, nitrate/nitrite antiporter | 0.019 | 0.015 |
| Hden_0926 | Nitrate reductase subunit alpha, NarG | 0.179 | 0.152 |
| Hden_1054 | Cytochrome c | 0.160 | 0.172 |
| Hden_1055 | Cytochrome $c_{550}$ domain protein | 0.007 | 0.010 |
| Hden_1483 | Cytochrome c family protein | 0.480 | 1.000 |
| Hden_1879 | Permease protein NosY, copper transport | 0.408 | 1.000 |
| Hden_1880 | ABC transporter ATP-binding protein NosF, copper transport | 0.466 | 0.460 |
| Hden_1881 | Periplasmic nitrous oxide reductase family maturation protein NosD | 0.266 | 0.306 |
| Hden_1882 | TAT-dependent nitrous-oxide reductase NosZ | 0.126 | 0.130 |
| Hden_1883 | NosR/NirI family protein | 0.031 | 0.037 |
| Hden_1884 | Ferritin family protein | 0.247 | 0.253 |
| Hden_1937 | NADH-quinone oxidoreductase subunit NuoF | 0.481 | 1.000 |
| Hden_2045 | FixH family protein | 0.107 | 0.107 |
| Hden_2046 | Cytochrome c oxidase accessory protein CcoG | 0.064 | 0.071 |
| Hden_2047 | Cytochrome-*c* oxidase *cbb*$_3$-type subunit III | 0.012 | 0.014 |
| Hden_2048 | cbb3-type cytochrome c oxidase subunit 3 | 0.006 | 0.010 |
| Hden_2049 | Cytochrome-*c* oxidase *cbb*$_3$-type subunit II | 0.007 | 0.011 |
| Hden_2050 | Cytochrome-*c* oxidase *cbb*$_3$-type subunit I | 0.005 | 0.008 |
| **Carbon metabolism** | | | |
| Hden_0042 | Poly(3-hydroxybutyrate) depolymerase | 0.435 | 0.484 |
| Hden_0607 | NAD-dependent formate dehydrogenase | 3.915 | 2.864 |
| **Sulfur metabolism** | | | |
| Hden_0759 | SufS family cysteine desulfurase | ns | 0.472 |
| Hden_1046 | Sulfate adenylyltransferase subunit CysN | 0.250 | 0.388 |
| Hden_1047 | Sulfate adenylyltransferase subunit CysD | 0.426 | ns |

| | | | |
|---|---|---|---|
| Hden_1491 | NADPH-dependent assimilatory sulfite reductase hemoprotein subunit, CysI | 0.433 | ns |
| **Transport** | | | |
| Hden_2042 | Sulfite exporter TauE/SafE family protein | 0.438 | 0.376 |
| Hden_2044 | cadmium-translocating P-type ATPase | 0.151 | 0.142 |
| Hden_2136 | DHA2 family efflux MFS transporter permease subunit | 0.384 | 0.261 |
| **Heme degradation and iron acquisition** | | | |
| Hden_0575 | FtrA, periplasmic iron binding protein | 0.014 | 0.019 |
| Hden_0576 | HemN, oxygen-independent coproporphyrinogen III oxidase | 0.040 | 0.043 |
| Hden_0599 | Heme anaerobic degradation, anaerobilin synthase ChuW/HutW | 0.239 | 0.185 |
| **Regulation** | | | |
| Hden_0099 | PAS domain-containing protein | 0.118 | 0.120 |
| Hden_2177 | Crp/Fnr family transcriptional regulator | 0.022 | 0.021 |
| Hden_2274 | NnrS family protein, involved in response to NO | 0.391 | 0.360 |
| Hden_3436 | response regulator | 0.441 | 1.000 |
| **Other** | | | |
| Hden_0086 | Group II truncated hemoglobin | 0.092 | 0.087 |
| Hden_0095 | HPF/RaiA family ribosome-associated protein | 0.070 | 0.085 |
| Hden_0096 | Zinc-dependent alcohol dehydrogenase family protein | 0.036 | 0.040 |
| Hden_0097 | Flavin reductase family protein | 0.070 | 0.074 |
| Hden_0174 | Hypothetical protein | 0.261 | 1.000 |
| Hden_0328 | Hypothetical protein | 0.399 | 0.361 |
| Hden_0672 | TonB-dependent receptor | 0.150 | 0.151 |
| Hden_0959 | 5-aminolevulinate synthase | 0.400 | 0.430 |
| Hden_1119 | Phage tail tape measure protein | ns | 0.444 |
| Hden_1171 | Hypothetical protein | 0.491 | ns |
| Hden_1773 | radical SAM protein | 0.272 | 0.313 |
| Hden_1841 | Universal stress protein | 0.099 | 0.102 |
| Hden_1876 | Hypothetical protein | 0.038 | 0.043 |
| Hden_2272 | Membrane protein | 0.276 | 0.180 |
| Hden_2281 | HD domain-containing protein | ns | 0.464 |
| Hden_2596 | Alpha/beta fold hydrolase | 0.395 | 0.367 |
| Hden_2615 | Hypothetical protein | 0.117 | 0.092 |
| Hden_2827 | Ferric reductase-like transmembrane domain-containing protein | 0.024 | 0.025 |
| Hden_2910 | Hypothetical protein | 0.397 | 0.458 |
| Hden_3135 | Circularly permuted type 2 ATP-grasp protein | 0.487 | ns |

ns, not significant

[a] Gene names obtained using sequence similarities in Uniprot or NCBI databases

** Significance threshold set at <0.5-fold change and $p$<0.001;

Simpson, A.J., Reinach, F.C., Arruda, P., Abreu, F.A., Acencio, M., Alvarenga, R., Alves, L.M., Araya, J.E., Baia, G.S., Baptista, C.S., Barros, M.H., Bonaccorsi, E.D., Bordin, S., Bove, J.M., Briones, M.R., Bueno, M.R., Camargo, A.A., Camargo, L.E., Carraro, D.M., Carrer, H., Colauto, N.B., Colombo, C., Costa, F.F., Costa, M.C., Costa-Neto, C.M., Coutinho, L.L., Cristofani, M., Dias-Neto, E., Docena, C., El-Dorry, H., Facincani, A.P., Ferreira, A.J., Ferreira, V.C., Ferro, J.A., Fraga, J.S., Franca, S.C., Franco, M.C., Frohme, M., Furlan, L.R., Garnier, M., Goldman, G.H., Goldman, M.H., Gomes, S.L., Gruber, A., Ho, P.L., Hoheisel, J.D., Junqueira, M.L., Kemper, E.L., Kitajima, J.P., Krieger, J.E., Kuramae, E.E., Laigret, F., Lambais, M.R., Leite, L.C., Lemos, E.G., Lemos, M.V., Lopes, S.A., Lopes, C.R., Machado, J.A., Machado, M.A., Madeira, A.M., Madeira, H.M., Marino, C.L., Marques, M.V., Martins, E.A., Martins, E.M., Matsukuma, A.Y., Menck, C.F., Miracca, E.C., Miyaki, C.Y., Monteriro-Vitorello, C.B., Moon, D.H., Nagai, M.A., Nascimento, A.L., Netto, L.E., Nhani, A., Jr., Nobrega, F.G., Nunes, L.R., Oliveira, M.A., de Oliveira, M.C., de Oliveira, R.C., Palmieri, D.A., Paris, A., Peixoto, B.R., Pereira, G.A., Pereira, H.A., Jr., Pesquero, J.B., Quaggio, R.B., Roberto, P.G., Rodrigues, V., de, M.R.A.J., de Rosa, V.E., Jr., de Sa, R.G., Santelli, R.V., Sawasaki, H.E., da Silva, A.C., da Silva, A.M., da Silva, F.R., da Silva, W.A., Jr., da Silveira, J.F., Silvestri, M.L., Siqueira, W.J., de Souza, A.A., de Souza, A.P., Terenzi, M.F., Truffi, D., Tsai, S.M., Tsuhako, M.H., Vallada, H., Van Sluys, M.A., Verjovski-Almeida, S., Vettore, A.L., Zago, M.A., Zatz, M., Meidanis, J., and Setubal, J.C. (2000) The genome sequence of the plant pathogen *Xylella fastidiosa*. The Xylella fastidiosa Consortium of the Organization for Nucleotide Sequencing and Analysis. *Nature* **406**: 151-159.

Tanabe, T.S., and Dahl, C. (2023) HMSS2: an advanced tool for the analysis of sulfur metabolism, including organosulfur compound transformation, in genome and metagenome assemblies. *Molecular Ecology Resources* **23**: 1930-1945.

Tatusova, T., Ciufo, S., Fedorov, B., O'Neill, K., and Tolstoy, I. (2014) RefSeq microbial genomes database: new representation and annotation strategy. *Nucleic Acids Research* **42**: D553-D559.

Thöny-Meyer, L. (2002) Cytochrome *c* maturation: a complex pathway for a simple task? *Biochemical Society Transactions* **30**: 633-638.

Valdes, J., Ossandon, F., Quatrini, R., Dopson, M., and Holmes, D.S. (2011) Draft genome sequence of the extremely acidophilic biomining bacterium *Acidithiobacillus thiooxidans* ATCC 19377 provides insights into the evolution of the *Acidithiobacillu*s genus. *Journal of Bacteriology* **193**: 7003-7004.

Venkatramanan, R., Prakash, O., Woyke, T., Chain, P., Goodwin, L.A., Watson, D., Brooks, S., Kostka, J.E., and Green, S.J. (2013) Genome sequences for three denitrifying bacterial strains isolated from a uranium- and nitrate-contaminated subsurface environment. *Genome Announcements* **1**.

Weinitschke, S., Denger, K., Cook, A.M., and Smits, T.H.M. (2007) The DUF81 protein TauE in *Cupriavidus necator* H16, a sulfite exporter in the metabolism of $C_2$ sulfonates. *Microbiology* **153**: 3055-3060.

Weissgerber, T., Zigann, R., Bruce, D., Chang, Y.J., Detter, J.C., Han, C., Hauser, L., Jeffries, C.D., Land, M., Munk, A.C., Tapia, R., and Dahl, C. (2011) Complete genome sequence of *Allochromatium vinosum* DSM 180$^T$. *Standards in Genomic Sciences* **5**: 311-330.

Williams, S.G., Attridge, S.R., and Manning, P.A. (1993) The transcriptional activator HlyU of *Vibrio cholerae*: nucleotide sequence and role in virulence gene expression. *Molecular Microbiology* **9**: 751-760.

Wodara, C., Bardischewsky, F., and Friedrich, C.G. (1997) Cloning and characterization of sulfite dehydrogenase, two *c*-type cytochromes, and a flavoprotein of *Paracoccus denitrificans* GB17: essential role of sulfite dehydrogenase in lithotrophic sulfur oxidation. *Journal of Bacteriology* **179**: 5014-5023.

Wu, X.L., Yu, S.L., Gu, J., Zhao, G.F., and Chi, C.Q. (2009) *Filomicrobium insigne* sp. nov., isolated from an oil-polluted saline soil. *International Journal of Systematic and Evolutionary Microbiology* **59**: 300-305.

Zhang, Y., Kitajima, M., Whittle, A.J., and Liu, W.T. (2017) Benefits of genomic insights and CRISPR-Cas signatures to monitor potential pathogens across drinking water production and distribution systems. *Frontiers in Microbiology* **8**: 2036.
